# Epigenetic memory acquired during long-term EMT induction governs the recovery to the epithelial state

**DOI:** 10.1101/2022.08.23.504814

**Authors:** Paras Jain, Sophia Corbo, Kulsoom Mohammad, Jason T George, Herbert Levine, Michael Toneff, Mohit Kumar Jolly

## Abstract

Epithelial-Mesenchymal Transition (EMT) and its reverse Mesenchymal-Epithelial Transition (MET) are critical during embryonic development, wound healing and cancer metastasis. While phenotypic changes during short-term EMT induction are reversible, long-term EMT induction has been often associated with irreversibility. Here, we show that phenotypic changes seen in MCF10A cells upon long-term EMT induction by TGFβ need not be irreversible, but have relatively longer timescales of reversibility than those seen in short-term induction. Next, using a phenomenological mathematical model incorporating the epigenetic silencing of miR-200 by ZEB, we highlight how the epigenetic memory gained during long-term EMT induction can slow the recovery to the epithelial state post-TGFβ withdrawal. Our results suggest that epigenetic modifiers can govern the extent and timescale of EMT reversibility, and advise caution against labelling phenotypic changes seen in long-term EMT induction as ‘irreversible’.

## Introduction

Cellular plasticity – the ability of cells to reversibly alter their phenotypes – is a hallmark of cancer metastasis (Welch and Hurst, 2019). One of the most well-investigated axes of cellular plasticity is epithelial-mesenchymal plasticity (EMP) which involves reversible transitions between epithelial (E), mesenchymal (M) and hybrid E/M phenotypes. Aside from metastasis, EMP is also implicated in embryonic development, wound healing, tissue repair and fibrosis (Yang et al., 2020). Initially thought of as a binary process, EMP is now understood to encompass many hybrid E/M states and incorporate reversible spontaneous or induced switching among these multiple phenotypes (Tripathi et al., 2020b). Studying the dynamics of EMP studied in cancer cell lines (patient or animal derived) have revealed a phenotypically heterogeneous distribution of E-M states, such that temporal evolution starting from different subpopulations can often recapitulate the parental heterogeneous distribution, given enough time (Bhatia et al., 2019b; Pastushenko et al., 2018; Ruscetti et al., 2016). The equilibrium distribution of E-M states in a cell population can be regulated by external conditions, including changes in the concentrations of growth factors in culture media and the presence of cytotoxic/cytostatic drugs (Gupta et al., 2011; Jagannathan et al., 2020; Ruscetti et al., 2016). Further, cell-cell communication through juxtacrine, autocrine and paracrine signalling can also influence cell-state switching and modulate phenotypic heterogeneity (Boareto et al., 2016; Neelakantan et al., 2017; Yamamoto et al., 2017). All these observations suggest dynamic interconversion among the E, M and hybrid E/M phenotypes.

At an intracellular level, EMP is enabled by a complex interplay of diverse molecules and signalling pathways involved in feedback loops. For instance, a mutual inhibitory feedback loop between the families of transcription factor ZEB and micro-RNA miR-200, and the positive feedback loop between ZEB and TGF-β signalling pathway are canonical mechanisms of EMP across multiple cancer types (Burk et al., 2008; Park et al., 2008; Tian et al., 2013; Watanabe et al., 2019). Mathematical modeling of these interactions has identified hysteresis on transitioning between E and M states, i.e. the concentration of an EMT inducing signal for which cells transition from an E to a M state become different from those utilized for reversion to an E state (Lu et al., 2013). Hysteretic dynamics has been confirmed in many *in vitro* studies since (Celià-Terrassa et al., 2018; Karacosta et al., 2019). The presence of hysteresis often implies that once cells attain a M state, it is difficult to regain an E state even if the external inducer (e.g. TGFβ) is removed/withdrawn from the culture media. However, recent studies report that while epithelial cells can attain a M state within a short term exposure to Epithelial Mesenchymal Transition (EMT) inducers, a threshold exposure time to these signals is required for a population to undergo an ‘effectively irreversible’ state change to attain a ‘stabilized EMT’ state (Jia et al., 2019; Katsuno et al., 2019). The extent of this irreversibility can vary depending on genetic background of the cell, induction factor and/or dose. For instance, long-term treatment of MCF10A cells with TGFβ induced chromatin accessibility changes among genes associated with EMT, apico-basal polarity and stemness. However, most cells also regained certain epithelial traits such as E-cad localization, morphology, and loss in migratory ability in the experimental time window (Johnson et al., 2022). Thus, further investigation is needed to better understand the dynamics of reversible EMT and MET *in vitro* and *in vivo*.

Epigenetic modifications such as DNA (de-)methylation and histone-level (de-)acetylation at various promoter regions can underlie such effectively ‘irreversible’ changes. When immortalized HMEC cells expressing oncogenic Ras were cultured in 10% serum for multiple passages, the promoter region of E-cadherin became increasingly methylated (Dumont et al., 2008). Similarly, in MDCK cells, autocrine TGFβ signalling maintained promoter methylation of micro-RNA 200b-c, contributing to a stabilized M state (Gregory et al., 2011). Inhibiting this autocrine signalling reduced the micro-RNA 200b-c promoter methylation levels and thus allowed a Mesenchymal-Epithelial Transition (MET). Consistently, overexpression of miR-200 together with the knockdown of chromatin remodelling protein BRG1 was required to induce a MET in RD sarcoma cells (Somarelli et al., 2016). Together, these reports suggest that chromatin reprogramming can control the reversible dynamics of EMT/MET. Besides epigenetic aspects, different modules of genes whose expression patterns are altered during induction of EMT can retreat to their pre-treatment levels at varying rates, as seen in prostate cancer cells (Stylianou et al., 2019). Such timescales differences in the reversibility of transcriptomic changes can lead to distinct EMT/MET trajectories at a single-cell level (Cook and Vanderhyden, 2020; Karacosta et al., 2019) and have been implicated in faster switching to a drug resistant state when cells are re-exposed to drugs after a short drug holiday (Pisco et al., 2013). However, it remains unclear whether the observed time-scale differences are due to differences in half-lives of mRNAs and/or proteins, slowly evolving and/or accumulating epigenetic modifications, or a combination of these effects.

Here, using experiment and mathematical modelling approaches, we show that the timescales of reversibility (MET) at the population level depends on the duration for which EMT was induced. We used TGFβ to induce EMT in MCF10A cells for two different durations (13 days, 22 days) and measured, at the population level, both gain in epithelial genes (miR-200b, miR-200c, and E-cad) expression and loss of mesenchymal gene (ZEB) expression, up to 18 days post-TGFβ withdrawal (for 13 days induction) and up to 45 days post-withdrawal (for 22 days induction). We hypothesized that the epigenetic repression of miR-200 by ZEB, and consequent accumulation of ‘epigenetic memory’ can prevent MET, thus explaining the timescale differences in reversibility as seen experimentally for short vs long treatment with TGFβ. We adopted our earlier mathematical modeling formalism capturing the epigenetic repression of miR-200 by ZEB in a phenomenological manner (Jia et al., 2019). Our model predicts that while prolonged treatment with an EMT inducer can lead to a slower MET due to differences in signaling activation levels, the accumulation of epigenetic memory appears to be the major determinant of the difference in reversal timescales between short- and long-term EMT induction. Thus, altering the rate of accumulation and/or decay of epigenetic memory, through treatment with various epigenetic modifiers, can govern the extent of reversibility of EMT. Further, our stochastic simulations demonstrate population heterogeneity at a single-cell level by quantifying the time taken to revert to an epithelial state post-withdrawal of the EMT-inducing signal. Overall, our analysis highlights how the timescale of EMT reversibility may depend on the duration of EMT induction and consequent epigenetic changes, and advises caution against mislabelling changes witnessed during long-term EMT induction as ‘irreversible’.

## Results

### TGFβ treated MCF10A cells for extended durations require longer withdrawal time to revert to an epithelial state

Earlier experiments in MCF10A cells suggest an effectively ‘irreversible’ switch to a mesenchymal state, by treatment with TGFβ for up to 15 days and withdrawal for 15 days post-treatment (Jia et al., 2019). To interrogate the dependence of EMT reversibility on TGFβ induction duration as well as that of withdrawal, we considered two different time-scales: a) short-term, i.e. exposure of MCF10A cells to TGFβ for 13 days and under observation for up to 18 days post-withdrawal, and b) long-term, i.e. exposure of MCF10A cells to TGFβ for 22 days and under observation for up to 45 days post-withdrawal (**Fig 1A**). We found that for the short-term treatment of 13 days, the expression level of epithelial genes (*CDH1, miR-200b, miR-200c*) and that of mesenchymal marker *ZEB1* returned to their pre-treatment levels within roughly 18 days of withdrawal (**Fig 1B-C**). However, the time taken to return to the pre-treatment level was much higher when the EMT induction consisted of 22 days of TGFβ exposure. With this protocol, it took almost 45 days after withdrawal to revert to pre-treatment levels (**Fig 1B-C**). Together, these preliminary observations clearly suggested that the timescale of MET – observed through canonical markers measured at the bulk level – depended strongly on the time-period of induction, and that MET need not be completely ‘irreversible’ as proposed earlier, even after chronic treatment.

**Fig. 1.**
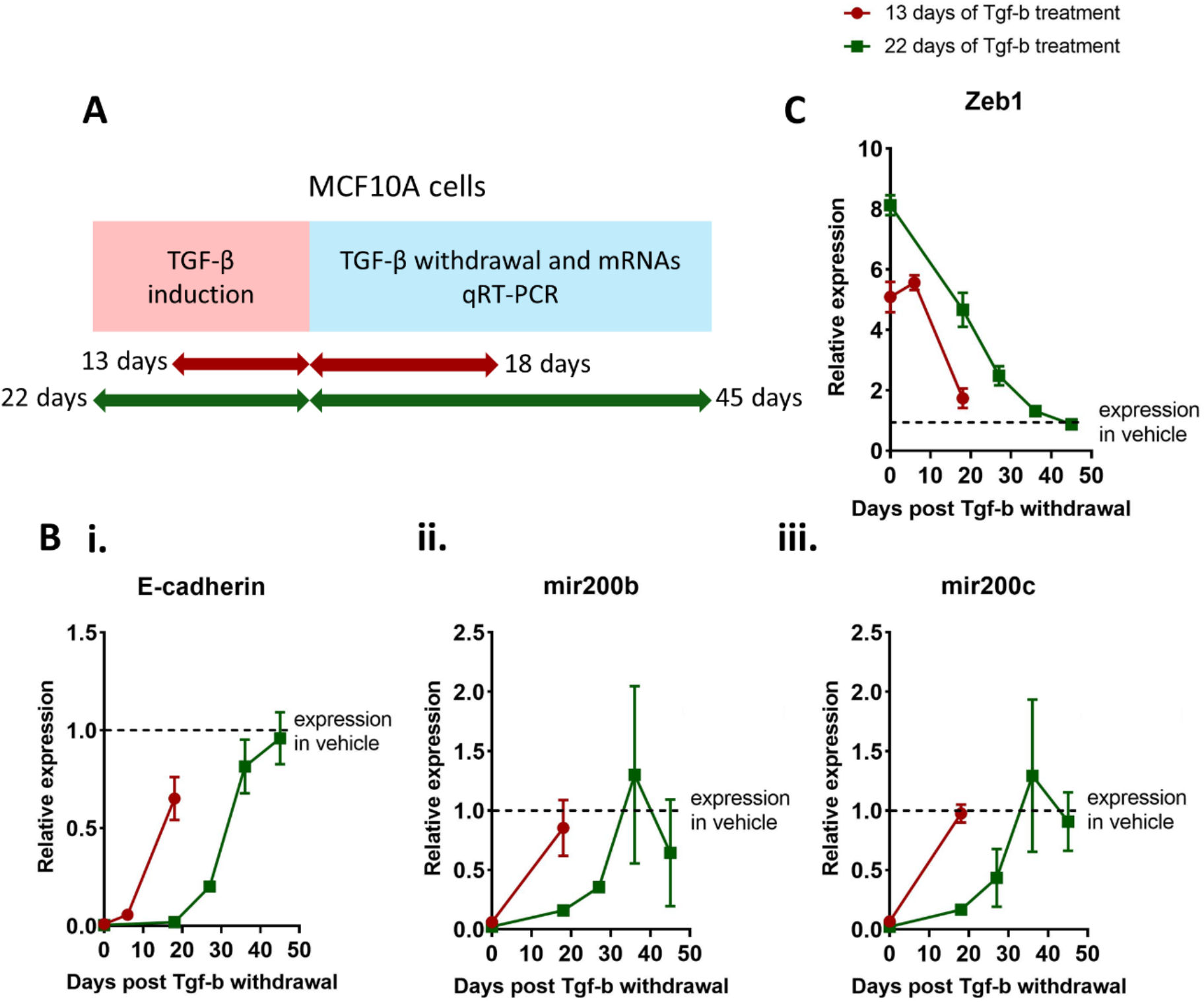
Time taken to regain basal expression of Epithelial and mesenchymal marker (at population level) after short (13 days) and long-term (22 days) EMT induction. **A)** Experimental strategy – MCF10A cells were treated for 13 and 22 days with 5ng/ml TGFβ. After removal of TGFβ, cells were cultured until basal expression levels of epithelial and mesenchymal genes, prior to EMT, were regained. **B-C)** Reversal of B i) E-cad, B ii) miR200c, B iii) miR200c, and C) ZEB1 to their basal expression levels (relative to vehicle control) after short and long-term EMT induction. Data shown is averaged over three independent replicates. For 13 days EMT induction, transcripts E-cad and ZEB1 transcripts were quantified at 0, 6, 18 days post-withdrawal and miR-200b-c transcripts were quantified at 0, 18 days post-withdrawal. For 22 days EMT induction, all transcripts above were quantified at 0, 18, 27, 36, 45 days post-withdrawal.

Next, we observed a marked difference in the temporal trajectories defining the recovery of expression levels for E-cadherin, ZEB1 and miR-200 under 22 day of TGFβ treatment. Here, E-cadherin levels did not increase for the first 18 days post-withdrawal, showing an initial silenced phase, but then exhibited sigmoidal increase trend over the next 27 days, before ultimately saturating (**Fig 1B, i**). Similar patterns showing an initial lag period were observed for miR-200b and miR-200c levels as well (**Fig 1B, ii-iii**). ZEB1 levels, on the other hand, followed near-linear decrease for the first 27 days and later plateaued (**Fig 1C)**. Together, these trends highlight two points: a) different genes may exhibit varying dynamics of recovery, reminiscent of similar observations in LNCaP cells (Stylianou et al., 2019), and b) caution should be exercised before claiming ‘irreversible’ EMT, depending on the timescales and the specific markers/features by which the reversal was assessed experimentally. For instance, at 18 days post-withdrawal, E-cadherin dynamics may suggest seemingly irreversible EMT, yet E-cadherin levels were restored around day 45 post-withdrawal (**Fig 1B, i**).

### Mathematical model of epigenetic regulation in Epithelial-Mesenchymal Transition

The abovementioned recovery dynamics of E-cadherin, miR-200b and miR-200c expression levels suggest a transient locking or stabilization of the mesenchymal state observed post-withdrawal (**Fig 1B**). These observations are reminiscent of the durable silencing of miR-200b and miR-200c expression observed in MDCK cells, post-TGFβ withdrawal, due to methylated promoter regions (Gregory et al., 2011). The extent of promoter methylation was shown to increase with the stabilization of mesenchymal state, caused either by the extended duration of TGFβ exposure or prolonged activation of self-sustaining autocrine loops such as those mediated between TGFβ signaling and ZEB1. Therefore, to better understand the role of epigenetic changes in mediating the recovery dynamics upon TGFβ exposure and withdrawal, we adapted our mechanism-based mathematical model of the EMT network (Tripathi et al., 2020b) to also incorporate epigenetic regulation. In our modeling framework, we consider that upon extended durations of treatment, ZEB1 can elicit methylation of the miR-200 promoter region (Davalos et al., 2012).

The regulatory network considered in our model includes interactions among miR-200, ZEB and SNAIL at transcriptional and translational levels (**Fig 2A**, inset) (Tripathi et al., 2020b). SNAIL represents the cumulative effects of TGFβ regulating EMT, and activates ZEB and inhibits miR-200. Further, ZEB and miR-200 inhibit each other, as experimentally reported (Burk et al., 2008; Park et al., 2008). In absence of any epigenetic regulation, the emergent dynamics of interactions among SNAIL, miR-200 and ZEB can allow for (co-)existence of multiple cell-states (phenotypes): mesenchymal (M; low miR-200, high ZEB), epithelial (E; high miR-200, low ZEB) and hybrid E/M (medium miR-200, medium ZEB) (**Fig 2A**; blue curves). In this bifurcation diagram, approximately, ZEB mRNA > 600 molecules correspond to a M state, ZEB mRNA < 150 molecules denote an E state, and ZEB mRNA between 150 and 600 molecules show a hybrid E/M state.

**Fig. 2.**
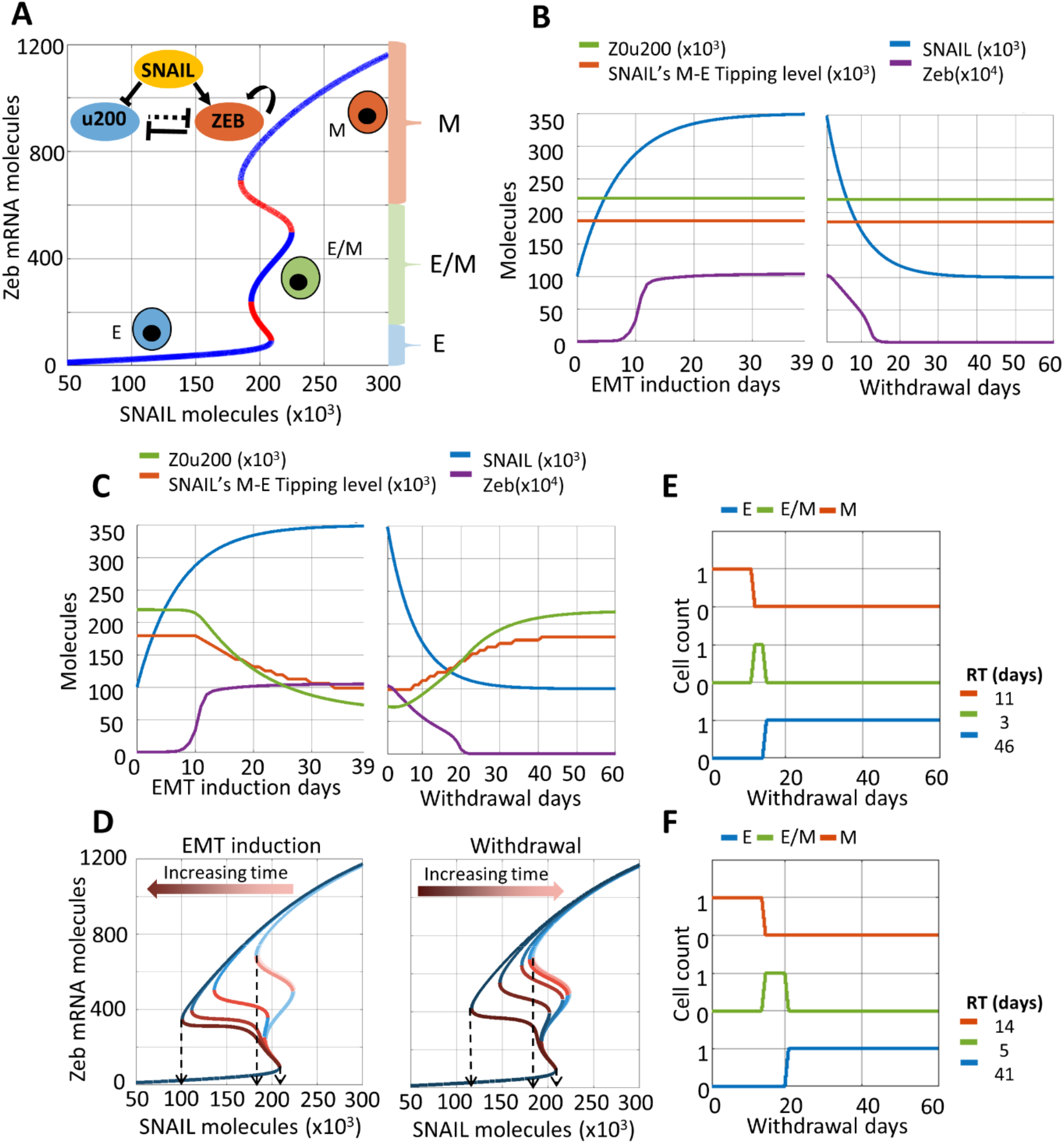
A phenomenological mathematical model to capture epigenetic regulation during EMT. **A)** (Inset) Regulatory network incorporating mutual inhibition between epithelial (mir200 – u200) and mesenchymal (ZEB) players, with SNAIL as external input. Bifurcation diagram shows equilibrium levels of ZEB mRNA based on SNAIL levels, as resulting from dynamics of the regulatory network. Blue curves are stable equilibria while red ones are unstable. Three distinct coloured braces on the right (red, green, and blue) qualitatively represents the ZEB mRNA levels used to assign epithelial (E), mesenchymal (M) or hybrid E/M phenotype. **B)** Dynamics of ZEB levels with changing SNAIL levels during EMT induction (left) and withdrawal (right) without considering epigenetic regulation of miR-200 by ZEB. **C)** Same as B) but with incorporating epigenetic regulation. Z0u200 levels and SNAIL M-E tipping levels reflect the extent of epigenetic reprogramming at any time instant. **D)** Bifurcation diagrams for ZEB mRNA levels for varying strengths of epigenetic regulation (Z0u200 levels) during EMT induction (left) or withdrawal (right) of EMT-inducing signal, at an interval of 10 days each (0, 10, 20, 30, 39 days post-induction, and 10, 20, 30, 40, 50, 60 days post-withdrawal). Black arrows highlight the SNAIL levels corresponding to E-to-M tipping (right arrow) and M-to-E tipping (left and middle arrows). **E)** Changes in cell’s phenotype during withdrawal period following EMT induction without epigenetic regulation. **F)** Same as E) but with the impact of epigenetic regulation. RT (residence time) measures the time that the cell spends in each phenotype during withdrawal period of 60 days. Parameters used in B and E: S_01_ = 100K molecules, S_02_ = 350K molecules, α = 0, β_for_ = 1 hr, and β_rev_ = 1 hr. When considering epigenetic regulation (panels C and F), α = 0.15, β_for_ = 240 hr, and β_rev_ = 240 hr.; all other parameters as abovementioned.

First, we investigate the dynamics of this network in absence of any epigenetic influence of ZEB on the miR-200 family promoters. In our simulation framework, the dynamics of SNAIL in a cell is modelled as a deterministic variable which tends to approach a saturation value (Eq. 2 in Methods, with zero noise amplitude), and we modulate the SNAIL saturating values to mimic EMT and MET induction in our model. Prior to EMT induction, a cell exhibits SNAIL levels corresponding to an E phenotype (approximately 100K molecules (**Fig 2A**)). During EMT induction, the cellular SNAIL level increases and eventually saturates at a much higher value (approximately 350K molecules), with a corresponding increase in ZEB levels (**Fig. 2B**, left), and acquisition of a mesenchymal state (**Fig 2A**). Upon withdrawal of the EMT inducing signal, the levels of SNAIL and ZEB gradually return to their initial values, thus reflecting MET (**Fig 2B**, right). Thus, in absence of any epigenetic regulation, our model could recapitulate the reversible EMT/MET dynamics.

Next, we examine how incorporating the epigenetic influence on miR-200 mediated by ZEB1 can alter the dynamics of EMT/MET. Experimental data, including ours (**Fig 1**), suggests that the longer a cell stays in the M state, the slower will be its reversibility dynamics following induction withdrawal (Dumont et al., 2008; Gregory et al., 2011). To obtain these dynamics, we assumed the threshold of ZEB levels needed to suppress miR-200 in the corresponding Hills function (Z0u200) to be a time-dependent function of ZEB levels (Eq. 1, Methods) (Jia et al., 2019). Thus, a higher saturating level of ZEB during EMT will continue to decrease the levels of Z0u200, enabling lower levels of ZEB to repress miR-200, and thereby incorporating the impact of epigenetic changes caused by ZEB (**Fig 2C**, left panel green curve vs. **Fig 2B**, left panel green curve). The dynamics of SNAIL and ZEB, however, remain unchanged during induction, as expected (compare corresponding blue and violet curves in **Fig 2C**, left panel vs. that in **Fig 2B**, left panel).

We further calculated how the temporally varying levels of Z0u200 during EMT induction and withdrawal altered the bifurcation diagram for the EMT network. As the levels of Z0u200 decreased during EMT induction, we saw no observable change in the tipping point levels of SNAIL required for cells to switch from the E to M state, but noticed a complimentary decrease in the tipping point for an M to E state switch (dotted arrows in **Fig 2D**). In other words, the epigenetic influence mediated by ZEB can reshape the phenotypic stability landscape such that it becomes more difficult for cells to revert to an epithelial state post-withdrawal. Such changes are not seen in the scenario where epigenetic changes are absent (compare orange curve in **Fig 2C**, left panel vs. that in **Fig 2B**, left panel). The longer the EMT induction period, the lower the Z0u200 levels; this trend can explain why short-term EMT induction is expected to have much weaker epigenetic impact as compared to long-term induction (**Fig 2D**). As the EMT-inducing signal is withdrawn, such accumulated epigenetic changes decay slowly, thus leading to recovery of Z0u200 molecules and SNAIL’s M to E tipping point to pre-induction levels (**Fig 2C, D**, right). This change ends the lag period in the recovery of levels of EMT/MET regulators (compare orange and green curves in **Fig 2C**, right panel vs. that in **Fig 2B**, right panel).

Finally, for comparative analysis, we calculated the recovery time for cells induced to undergo EMT with epigenetic changes vs. those without any such changes. We quantified the number of days for which ZEB levels are in the above-mentioned numerical ranges corresponding to the E, M and hybrid E/M states (**Fig 2A**). For the case without any epigenetic changes, the cells revert to an E state 14 days (= 11 days in the M state, followed by 3 days in the hybrid E/M state) days post-withdrawal (**Fig 2E**). However, when incorporating epigenetic influence, the cells stay in M and hybrid E/M states longer and return to an E state after 19 days (= 14 days in the M state, followed by 5 days in the hybrid E/M state), thus causing delayed recovery, or in other words, a slower MET (**Fig 2F**). The slower the decay of ‘epigenetic memory’ (Bintu et al., 2016; Hathaway et al., 2012) thus accumulated, the higher the delay in cells reverting to an epithelial state post-withdrawal of EMT-inducing signals.

### Longer EMT induction can delay the reversal to an epithelial state post-withdrawal of an EMT-inducing signal

Next, we interrogated how the duration of EMT induction can influence the build-up of ‘epigenetic memory’ and consequently the timescales of reversal to an epithelial state post-withdrawal of the EMT-inducing signal. We consider the response of a cell that is switching between two different values of SNAIL: S_01_ (prior to EMT induction) and S_02_ (post EMT-induction; thus S_02_ > S_01_). The timescale of epigenetic changes during induction or withdrawal is denoted by the rate of change of Z0u200 levels, given by β_for_ and β_rev_ respectively. The higher the value of β_for_, the slower the reduction in Z0u200 levels and thus the slower the build-up of epigenetic memory. The higher the value of β_rev_, the slower the return to pre-induction Z0u200 levels and thus the slower the decay of epigenetic memory. We simulated the dynamics of a cell exposed to short-term (13 days) and long-term (39 days) duration of EMT induction, and quantified the reversal time in these two scenarios.

For both short-term and long-term induction cases, EMT was induced by changing SNAIL levels from S_01_ = 100K molecules to S_02_ = 350K molecules, with epigenetic regulation timescales taken as β_for_ = 240 hr, and β_rev_ = 720 hr. Although ZEB levels increased and then saturated around 15 days of induction, a longer-term induction led to lower Z0u200 levels as compared to short-term induction. Thus, a longer induction conferred a relatively stronger epigenetic memory, as denoted by changes in bifurcation diagrams (**Fig S1**) and thus in tipping points levels of SNAIL required for cells to switch from M to E state (**Fig 3A** vs **Fig 3B**; left panels). Consequently, upon withdrawal of the EMT-inducing signal (i.e. reducing SNAIL levels from S_02_ = 350K molecules to S_01_ = 100K molecules), the time taken to recover the levels of ZEB and Z0u200 to pre-induction values is slower for long-term induction as compared to short-term induction (**Fig 3A** vs **Fig 3B**; right panels).

**Fig. 3.**
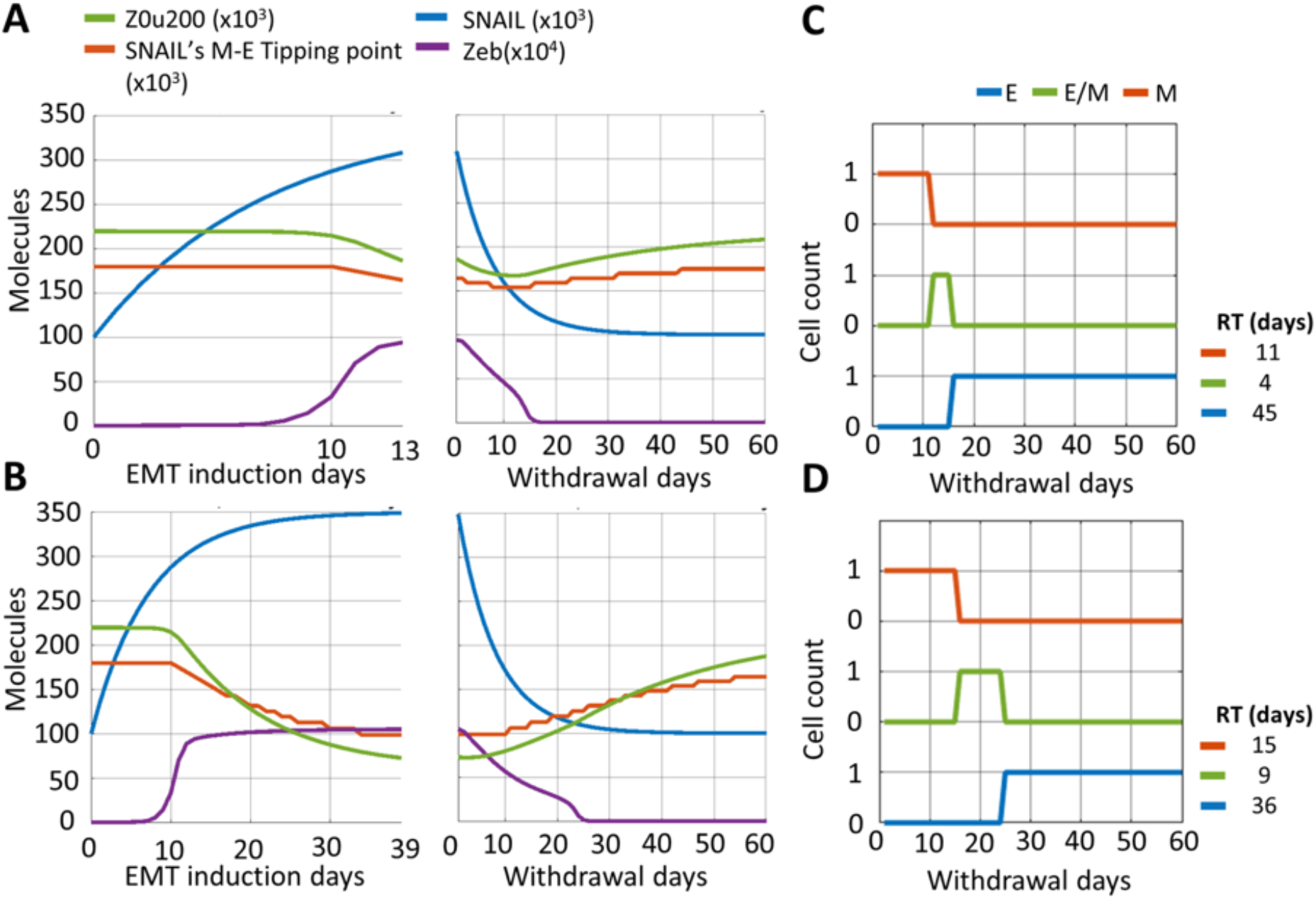
Influence of EMT induction time on recovery to an epithelial change post-withdrawal. Simulations for EMT induction for **A)** 13 days (short-term), and **B)** 39 days (long-term) followed by a withdrawal period of 60 days. Dynamics of ZEB, Z0u200 and the SNAIL levels corresponding to M-to-E tipping point are shown. **C, D)** Change in cell’s phenotype during withdrawal period after EMT induction for 13 days (A) and 39 days (B), respectively. RT (residence time) measures the time that the cell spends in each phenotype during withdrawal period of 60 days. Parameters used in panels A-D: α = 0.15, S_01_ = 100K molecules, S_02_ = 350K molecules, β_for_ = 240 hr, and β_rev_ = 720 hr.

A quantitative comparative analysis showed the difference in recovery time for cells induced to undergo EMT for different durations. For short-term induction, the cells revert to an E state 15 days (= 11 days in the M state, followed by 4 days in the hybrid E/M state) days post-withdrawal (**Fig 3C**). However, for long-term induction, this return happens after 24 days (= 15 days in the M state, followed by 9 days in the hybrid E/M one), possibly prolonging the residence of cells in hybrid E/M phenotype(s) (**Fig 3D**). These differences are much less prominent if the epigenetic feedback is not considered (**Fig S2**), thus showcasing the impact of a long-term EMT induction on epigenetic-level reprogramming.

### The rate of epigenetic changes during EMT induction and withdrawal determines the time taken to regain an epithelial cell state

The rate and extent of epigenetic reprogramming can depend on multiple factors, such as whether epigenetic changes are mediated via DNA methylation or by histone modification (Bintu et al., 2016). For instance, GRHL2, a canonical MET inducer, is a pioneering transcription factor capable of directly binding to condensed chromatin to initiate its opening, leading to cell-state changes (Balsalobre and Drouin, 2022; Chen et al., 2018). Such diverse modes of epigenetic regulation can alter the rate of accumulation (β_for_) and decay (β_rev_) of epigenetic memory.

We modulated the values of β_for_ and β_rev_ to assess their impact on epigenetic reprogramming and the timescales of cell-state transitions. First, we varied β_for_ (2x = doubled, 0.5x = halved) while maintaining β_rev_ = 720 hours. Reduced β_for_ values (= 120 hours) enhanced the rate of decrease of Z0u200 levels (**Fig S3A**), leading to a lower SNAIL’s M to E tipping levels at the end of induction period, as compared to the case for larger β_for_ values (= 480 hours) (**Fig 4A**). Higher β_for_ values result in lower accumulated epigenetic memory and thus, a faster reversal to an epithelial state post-withdrawal, irrespective of the induction period (**Fig 4B, S3B**). However, the epigenetic memory also saturates to a maximal level, as evident for smaller β_for_ values (**Fig 4B**). Second, we varied β_rev_ values while maintaining β_for_ = 240 hours. Larger β_rev_ values (= 1200 hours) led to a slower recovery from the acquired epigenetic memory accumulated during induction (**Fig S4A**), compared to smaller β_rev_ values (= 240 hrs) (**Fig 4C**), as witnessed by a difference in the slopes of curves of SNAIL’s M-E tipping level. The higher the value of β_rev_, the slower the decay of the accumulated epigenetic memory and thus, the longer the delay in reversal to an epithelial state post-withdrawal, irrespective of the induction period (**Fig 4D, S4B**).

**Fig. 4.**
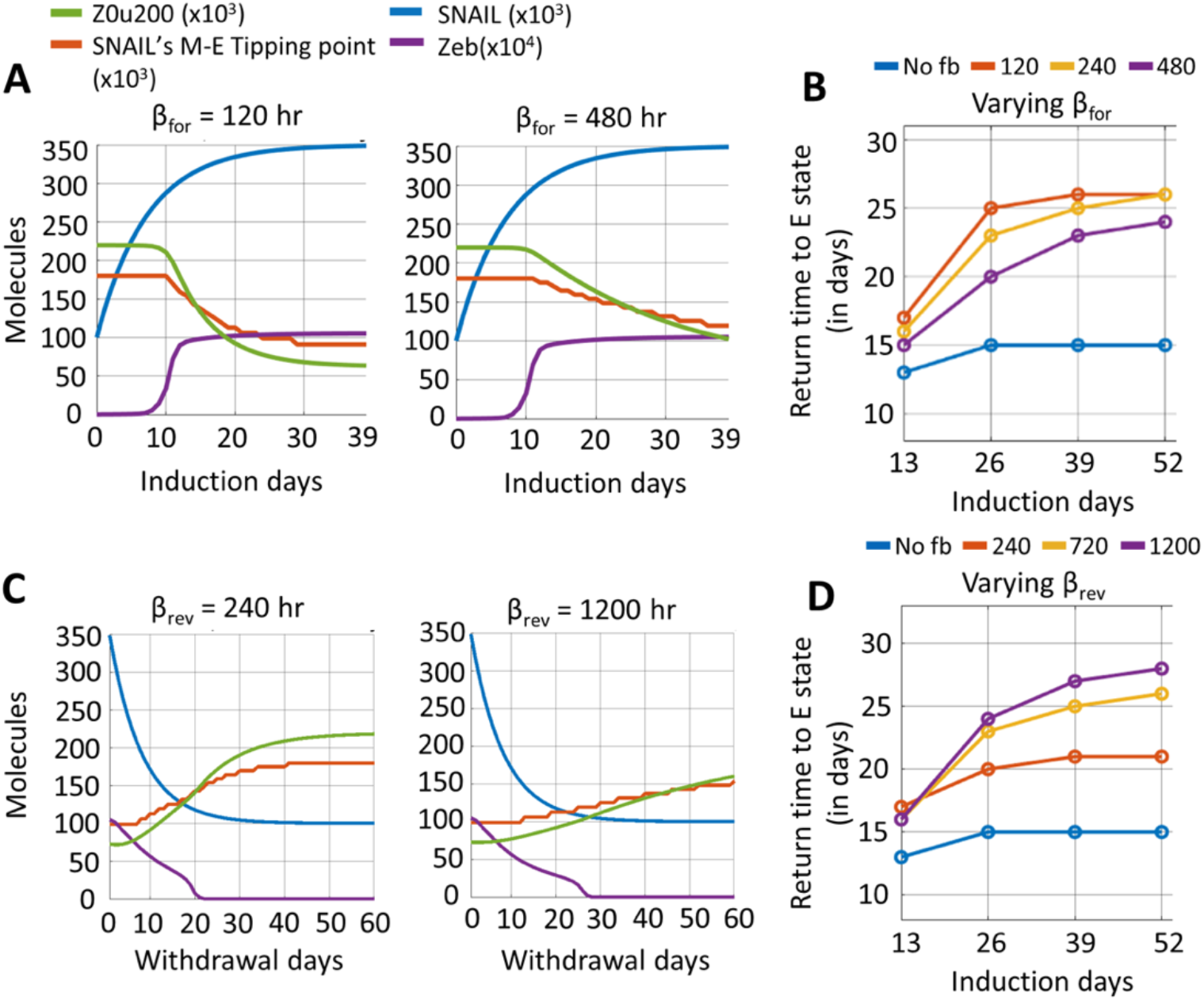
Effect of varying rates of acquisition and decay of epigenetic memory. **A)** Dynamics of ZEB, and changes in epigenetic regulation measures (Z0u200 and SNAIL levels corresponding to M-E tipping point) during EMT induction with variations in time constant β_for_ (acquisition rate of epigenetic changes, in units of hr), **B)** Time to recover to an E state for varied values of β_for_ and duration of EMT induction. **C)** Same as A) but for varied values of β_rev_ (timescale of decay of epigenetic changes in units of hr). **D)** Same as B) but for varied values of β_rev_ and duration of EMT induction. Parameter values, unless specified differently, are: α = 0.15, S_01_ = 100K molecules, S_02_ = 350K molecules, β_for_ = 240 hours, and β_rev_ = 720 hours., For cases without epigenetic regulation (no fb – no feedback), α = 0.

Together, these observations suggest that slower response at epigenetic regulation level, either during EMT induction or withdrawal, introduces a latency period for a cell to gain or lose epigenetic memory, thus impacting the rates of cell-state switching.

### Recovery timescales also depend on pre- and post-induction SNAIL levels

The extent of EMT/MET induction in a given cell can depend on multiple factors. These include the dose and duration of the inducing signal, pathways activated by the specific EMT/MET-inducing signal, and variations in the initial cell-state in terms of protein abundance or epigenetic status (Cook and Vanderhyden, 2020; Somarelli et al., 2016). To represent the impact of these factors, we examined the effects of varying the pre-induction (S_01_) and post-induction (S_02_) SNAIL levels on the time taken to recover to epithelial state post-withdrawal. We consider these scenarios both in the presence and absence of epigenetic feedback or memory.

In the absence of epigenetic regulation of miR-200 by ZEB1, we first examined the impact of varying S_01_. The lower the levels of S_01_, the faster the recovery dynamics. For S_01_= 100K molecules, for 39 days of EMT induction, it takes 14 days (11 days in the mesenchymal state followed by 3 days in the hybrid E/M state) to regain an epithelial phenotype (**Fig 2B, 2E**). However, at S_01_= 135K molecules, the recovery period extends to 18 days, and at S_01_= 170K molecules, it extends to 30 days (**Fig S5**). These slower dynamics of SNAIL can be attributed to the difference between post-induction values of SNAIL (S_02_) and the post-withdrawal values (S_01_) ones (**Eq. 2)**. However, on varying S_02_, the impact on recovery times is rather small, which can be explained by higher equilibrium levels of SNAIL achieved at the end of the EMT induction period. For S_02_= 300K molecules, it takes 12 days to revert to an epithelial state but for S_02_=400K molecules, it increases to 15 days (**Fig S6**). Thus, varying S_01_ levels had a stronger impact on the timescales of recovery to an epithelial state, than variations in S_02_ levels.

Next, we incorporated the epigenetic regulation of miR200 by ZEB and considered the case of a high S_01_ value (=135K molecules). The build-up of epigenetic memory, as seen before, delayed the time taken for SNAIL’s M-E tipping level to increase above the cellular SNAIL levels, and to attain pre-induction values during the period of withdrawal (**Fig 5A, S7A**). Consequently, the cell spent more time in M and E/M states and showed delayed recovery dynamics to an epithelial state. At even higher values of S_01_ (=170K molecules), the recovery slows further, eventually tending toward the scenario of irreversible EMT (**Fig 5B, S7B**).

**Fig 5.**
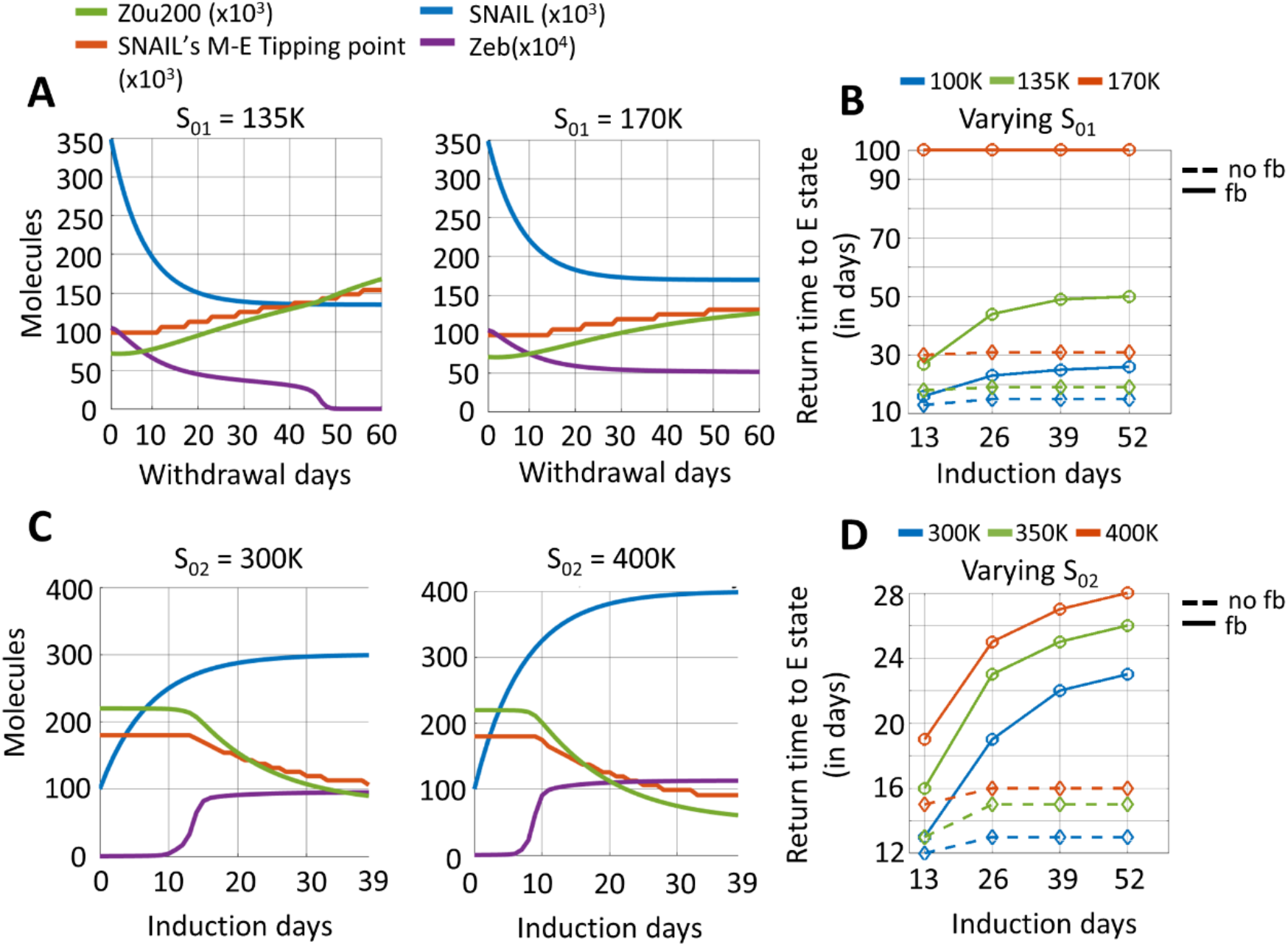
Effects of pre-induction (post withdrawal, S_01_) and post-induction (S_02_) SNAIL levels on epithelial recovery time following EMT induction. **A)** Dynamics of ZEB, and changes in epigenetic regulation measures – Z0u200 and SNAIL’s M-E tipping levels – during withdrawal period with variation in saturating basal SNAIL levels ‘S_01_’ with epigenetic changes during EMT. EMT induction duration was for 39 days. **B)** Time to recover epithelial state with variations in both S_01_ and EMT induction duration with epigenetic regulation (‘fb’, solid lines) and without epigenetic regulation (‘no fb’, dashed lines). **C)** Same as A but for varied saturating EMT-induced SNAIL levels ‘S_02_’ with epigenetic changes during EMT. **D)** Same as C but for varied S_02_ and EMT induction duration with epigenetic regulation (solid lines) and without it (dashed lines). Parameter values used, unless otherwise specified, are: S_01_ = 100K molecules, S_02_ = 350K molecules, β_for_ = 240 hr, β_rev_ = 720 hr, and α = 0.15.

Lastly, we varied the post-induction SNAIL levels (S_02_) while considering epigenetic changes. With increasing S_02,_ we observed higher steady levels of ZEB mRNA and protein. Also, increasing S_02_ accelerated EMT, thus stabilising ZEB in a high state for longer times during the induction period (**Fig 5C**). This prolonged time in a mesenchymal state during induction increased the gap between the tipping point levels of SNAIL for MET and EMT (**Fig S8**), thus acting as a barrier for MET. As expected, the higher the induction time, the stronger the extent of epigenetic memory accumulated, and thus the longer the time required to revert EMT (**Fig 5D**).

### Population-level effects of epigenetic changes during EMT

So far, we have examined the influence of epigenetic changes on reversibility towards an epithelial state, through simulating individual cells switching between two discrete levels of SNAIL (S_01_, S_02_). These simulations did not consider any stochastic fluctuations in protein levels. However, fluctuations in protein abundance can prevail at the single-cell level (Li et al., 2016) due to factors such as stochastic gene expression (Balázsi et al., 2011; Zhao et al., 2021) and asymmetry in cell division (Huh and Paulsson, 2011). To incorporate these factors, we model stochastic fluctuations around the mean level using the Ornstein-Uhlenbeck (OU) process. The OU process determines SNAIL levels by integrating a stochastic differential equation with both deterministic (drift) and stochastic (diffusion) terms (Eq. 2). Numerical implementation of the OU process provides SNAIL trajectories reflecting distinct cells that are statistically independent within the population (**Fig 6A**). The stochastic fluctuations in SNAIL levels were parameterised based on experimentally estimated values of the coefficient of variation (CV) of distributions of protein levels in a cellular population, and the 50% decorrelation time of single-cell expression levels (**Fig 6B, C**) (Li et al., 2016; Sigal et al., 2006).

**Fig 6.**
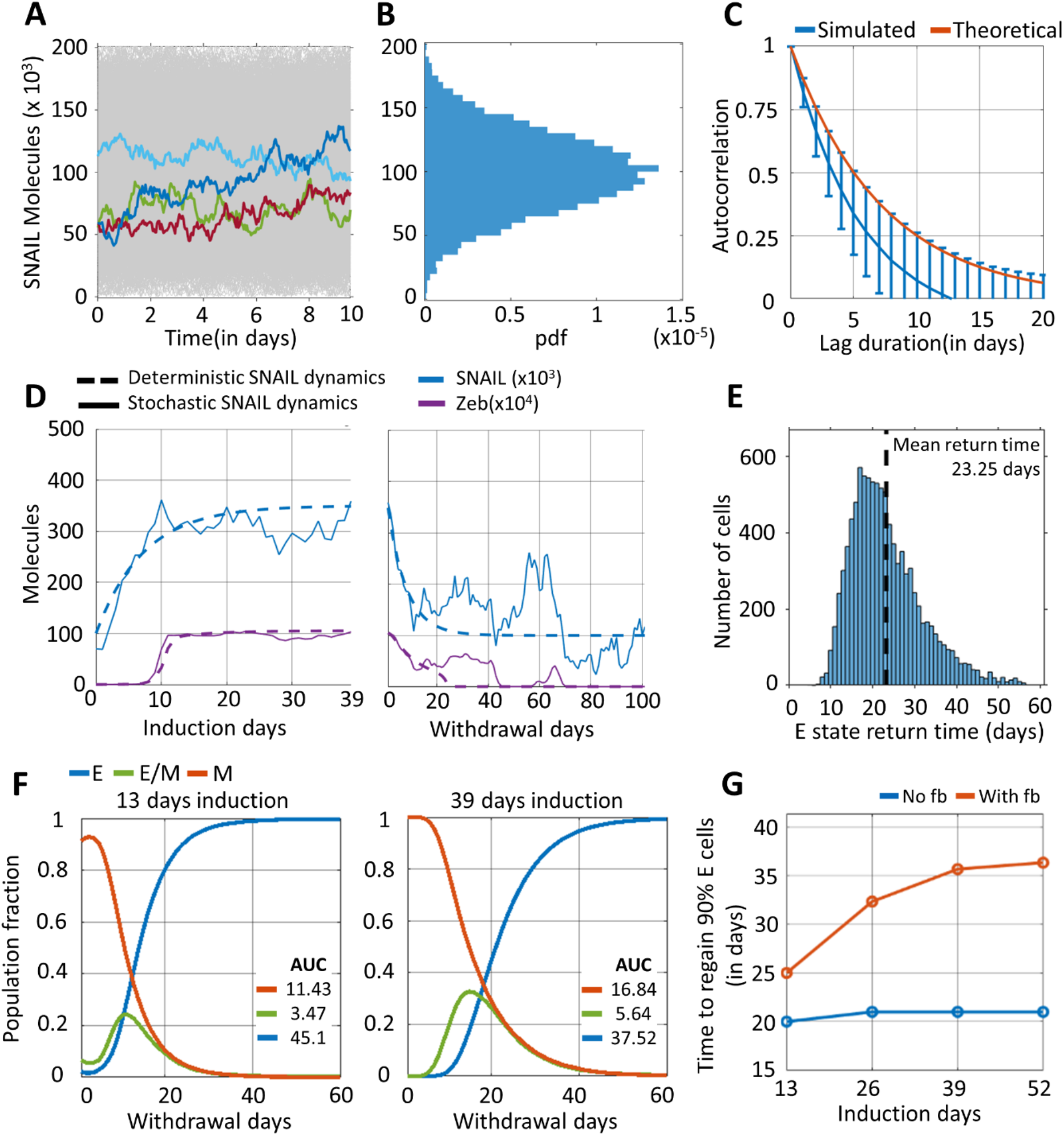
Effects of epigenetic changes during EMT on its reversibility in a heterogeneous population. **A)** Stochastic fluctuation in SNAIL levels around the population mean (S_0_ = 100K molecules). Temporal fluctuations in SNAIL levels derived from the total population of 10,000 cells are shown (in grey), of which 4 stochastic trajectories are coloured. **B)** The distribution of aggregate SNAIL levels at any realized time follows a Gaussian distribution, pdf refers to probability density function. **C)** Loss of correlation in SNAIL levels of a cell between any two time instants with increasing time separation. Red curve shows theoretical estimation of 50% decorrelation time as 5 days. Blue curve shows mean and standard deviation of decreasing autocorrelation of 10,000 independent fluctuating SNAIL time-series, each being 100 days long. **D)** Changes in SNAIL and ZEB levels during EMT induction and withdrawal period for a cell in the population, with and without stochastic fluctuations in SNAIL, shown by solid and dashed curves, respectively. **E)** Distribution of return time of cells to epithelial state after undergoing EMT induction for 39 days. Mean return time of population = 23.25 days (dashed line). **F)** Change in phenotypic distribution of population after undergoing EMT induction for 13 days and 39 days. ‘AUC’ corresponds to Area Under the Curve and reflects the cumulative fractional share of a phenotype in the population during the withdrawal period of 60 days. The results presented include the mean of three independent simulation runs of 10K cells each. **G)** Time taken to regain 90% of epithelial phenotype share in the population for increasing durations of EMT induction. Analysis was performed for scenarios with and without considering epigenetic changes during EMT, denoted by ‘with fb’ and ‘no fb’ respectively. Plots A-C were generated from analysing SNAIL time-series data. The time-series was stimulated using a stochastic differential equation (Eqn. 2). Plots D-G were obtained using following parameters for simulations: α = 0.15, S_01_ = 100K molecules, S_02_ = 350K molecules, β_for_ = 240 hr, and β_rev_ = 720 hr, for every cell in the population (in all simulated cases, the population size = 10,000 cells).

The stochastic simulations revealed heterogeneity in the dynamics of SNAIL and ZEB amongst individual cells within a population during EMT induction, and in the time taken to revert to an epithelial state upon withdrawal (**Fig 6D, E**; **Fig S9, S10**). Despite this heterogeneity, the mean return time (RT) to an epithelial state for a cell in the population remained close to our earlier observations from the deterministic analysis for a 39-day induction period. This similarity was seen for both scenarios – with the accumulation of epigenetic memory (mean RT = 23.25 days in **Fig 6E**, RT = 25 days in **Fig 4B**) as well as without it (mean RT = 14.9 days in **Fig S10B**, RT = 15 days in **Fig 4B**).

Next, we quantified the recovery dynamics to an epithelial phenotype, when a cell population has undergone EMT for varying durations, with and without epigenetic regulation of miR-200 by ZEB. We used two metrics – a) the cumulative fractional share of a phenotype in the population during the 60-day withdrawal period (**Fig 6F**), estimated by area under the curve (AUC) of phenotypic distribution plot, and b) the time taken for epithelial phenotype to comprise 90% of the population (**Fig 6G**). Without considering any epigenetic changes during EMT, a short-term (13 days) induction led to a faster recovery than a long-term (26 days and beyond) one (compare blue curves in **Fig S11** vs. in **Fig 6F**), reminiscent of earlier observations from the deterministic simulations (**Fig S1**). Upon accounting for epigenetic changes during EMT, the differences in recovery times became more pronounced with an increasing induction duration, suggesting a longer residence time of cells in M and hybrid E/M states (**Fig 6F** – compare AUC in left and right panels) and longer delays in regaining 90% epithelial share in the population (**Fig 6G**, red curve).

To ascertain how the rate of epigenetic changes during EMT and the withdrawal period influence the reversibility of EMT at a population level, we varied β_for_ and β_rev_ for every cell in the population. Increasing β_for_ lowered the residence time of cells in M and E/M states during withdrawal, thus speeding up the recovery to 90% E share (**Fig S12A-B**). Conversely, increasing β_rev_ enhanced the residence time of cells in M and E/M states during withdrawal, and delayed 90% E share recovery (**Fig S12C-D**). These results corroborated our observations during deterministic analysis (**Fig 4**). Similarities between single-cell and population-level analyses were also seen for varying S_01_ and S_02_ levels, where the changes in S_01_ had a more discernible impact on the dynamics of recovery than variations in S_02_ (for S_01_ variation, compare **Fig S5** with **Fig S13A, C** and **Fig 5B, Fig S7** with **Fig S13B**,**C**; for S_02_ variation, compare **Fig S6** with **Fig S14A, C** and **Fig 5B, Fig S8** with **Fig S14B**,**C**). Intriguingly and in contrast to the irreversible M state for higher S_01_ values (170K, **Fig 5A – right panel, Fig 5D**) observed for deterministic SNAIL dynamics, stochasticity in SNAIL levels enabled both a) cell transitions to the E state in the event of considerable dip in cellular SNAIL levels below M-E tipping levels (**Fig S15**), and b) spontaneous cell-state switching (**Fig S16**).

Overall, the dynamics of recovery to an epithelial state seen for deterministic SNAIL dynamics at individual cellular level were recapitulated by stochastic simulations for a cellular population whose SNAIL levels fluctuated around and switched between pre-defined mean SNAIL levels.

## Discussion

The co-existence of multiple cell-states along the E-M spectrum can be seen as “attractors” or valleys in a gene expression landscape, connected by trajectories that enable cell-state transitions (Burkhardt et al., 2022; Huang et al., 2009). The degree of resolution among distinct cell-states depends on the number of biomarkers used experimentally to identify a cell population (Bhatia et al., 2019a; Karacosta et al., 2019; Pastushenko et al., 2018; Pillai and Jolly, 2021; Ruscetti et al., 2016). These cell-states can transition between each other either spontaneously due to factors such as stochastic gene expression and asymmetric cell division, or under microenvironmental influence such as TGFβ signaling and altered matrix stiffness (Cook and Vanderhyden, 2020; Jain et al., 2022; Matte et al., 2019; Tripathi et al., 2020a; Zhao et al., 2021). The relative rates of cell-state transitions define the population distribution of cells along the E-M axis, as evident from dynamics of isolated subpopulations *in vitro* and *in vivo* (Bhatia et al., 2019b; Pastushenko et al., 2018; Yamamoto et al., 2017). These rates of transitions, and thus the equilibrium state distribution, are determined by the relative stability of each state (Hari et al., 2021).

Besides transcriptional and translational control, epigenetic regulation of E-M states can influence their relative stability, thus shaping population distributions (Brown et al., 2022; Tam and Weinberg, 2013). For instance, the hybrid E/M (EpCAM+ Vim+) and mesenchymal-like (EpCAM-Vim+) cells in the PKV cell line displayed upregulated levels of HMGA2, an epigenetic regulator (Ruscetti et al., 2016). Inhibiting HMGA2 using HDACi (Panobinostat) reduced the mesenchymal fraction of the population. Similarly, ectopic expression of EMT-TFs (SNAIL1, SNAIL2, ZEB1) in MDCK cells conferred a mesenchymal phenotype that included epigenetic silencing of the miR-200 family through DNA methylation (Diaz-Martin et al., 2014). While suppressing endogenous ZEB1 in cells ectopically expressing SNAIL1 did not revert cells to an epithelial state, SNAIL1 repression led to demethylation of the *miR-200b* and *miR-200c* genes, causing MET. In another context, ZEB1 recruited the epigenetic remodelling enzyme BRG1 at the promoter of E-cadherin and this regulation served as a barrier preventing GRHL2 from inducing MET (Somarelli et al., 2016). Thus, different EMT/MET-TFs can epigenetically control cellular plasticity. Consequently, combinatorial or sequential treatment with epigenetic regulators can govern the patterns of intratumor heterogeneity during metastasis and/or drug treatment (Ruscetti et al., 2016; Sharma et al., 2010).

Our mathematical model, which captures epigenetic changes during EMT, explains how epigenetic memory can accumulate as a function of the duration of an EMT-inducing signal, and how the reversibility of EMT depends on the rate of decay of this memory and thus the timepoint of withdrawal at which reversibility is experimentally assessed (Dumont et al., 2008; Gregory et al., 2011; Johnson et al., 2022). For instance, the loss in chromatin accessibility of the *EpCAM* gene was observed upon treatment of MCF10A cells with TGFβ showed recovery to pre-treatment levels for short-term treatment (4 days) but not for long-term treatment (10 days) (Johnson et al., 2022). Irreversible chromatin accessibility of many epithelial genes was demonstrated, but the withdrawal period was for 10 days only. Our data suggest that epithelial gene expression can be recovered after long-term EMT induction upon extended withdrawal periods (**Fig 1**), and this is likely reflected in the chromatin state of the cell. Therefore, it is possible that a re-opening of the chromatin state for *EPCAM* and other epithelial genes would have occurred over longer withdrawal times in the aforementioned study. Different timescales of cell-state reversal can be attributed to the dynamics of heterochromatin. For instance, in embryonic stem cells, a long-term recruitment (4.5 weeks) of heterochromatin protein 1 (HP1) to the *Oct4* promoter accumulated both H3K9me3 and DNA methylation that silenced Oct4 expression for multiple generations. However, a short-term recruitment (7 days) accumulated only H3K9me3 did not silence *Oct4* gene expression for long (Hathaway et al., 2012). Consistently, in pluripotent stem cells, the ratio of rate of methylation by DNMTs to the rate of demethylation Nanog-Tet complex determines the stability of the epigenetically silenced Oct gene (Chen et al., 2021). Similar differences in timescales of acquiring histone modification and DNA methylation was seen in the promoter of E-cadherin in HMLE cells grown in 10% serum condition (Dumont et al., 2008).

In our model formalism, the parameters β_for_ and β_rev_ capture distinct response times resulting from a variety of possible epigenetic regulators to reversibly and/or irreversibly silence a gene. While direct empirical identification of these individual rates is complicated by the fact that diverse epigenetic regulators often act in concert, synthetic biology approaches may be helpful in dissecting the dynamics of epigenetic regulation through different modifications: DNA methylation, histone deacetylation and histone methylation. For instance, in CHO-K1 cells, a synthetic genetic circuit was constructed to recruit histone methyl-transferase (EED, KRAB), histone deacetylase (HDAC4), and DNA methyltransferase (DNMT3B) to the fluorescent reporter gene (Bintu et al., 2016). These diverse regulators caused varied histone modification, with only DNMT3B recruitment leading to methylation at the promoter region. The distribution of gene silencing time at a single-cell level were quite distinct among these epigenetic modifiers after 80 hours of recruitment, with silencing due to DNMT3B being the slowest of all epigenetic regulators. While DNMT3B-mediated silencing did not lead to re-activation of expression during 30 days of observation time post-withdrawal, considerable recovery was observed for EEB, KRAB recruitment and full recovery to pre-treatment levels for HDAC4-mediated silencing. Further, increasing the duration of EED, KRAB and HDAC4 recruitment enhanced the fraction of cells showing silenced expression in the population after 30 days of withdrawal (Bintu et al., 2016). This observation corroborates with our findings that the extent and durability of epigenetic changes depends on duration of epigenetic modifier recruitment (in our case, maintenance of high ZEB levels during EMT, and its recruited epigenetic modifiers). It is conceivable that we can capture both the short and long-term memory effects of epigenetic changes by allowing a cell transition between three gene expression states: a) active gene expression state, b) reversibly silent state, and c) irreversibly silent gene state, such that the transition rates among these states dependent on the dose and/or duration of a specific epigenetic modifier treatment (Bintu et al., 2016; Mukund and Bintu, 2022).

Another factor that can vary the cellular response to external exposure to an EMT inducer (such as TGFβ) is cell-to-cell variability in protein levels. For instance, variation in the concentrations of TGFβ receptor and SMAD TFs among cells dictated their response to TGFβ stimulation in terms of nuclear localization of SMAD2 protein (Strasen et al., 2018). The cells were clustered into six response classes, whose proportions in the population varied with the concentration of TGFβ treatment. Similarly, cellular variability in levels of multiple proteins (DR4/5 receptors, DISC components, CASP8 and BID) controlled the time of apoptotic event in HeLa cells, in the case of TRAIL induced apoptosis (Spencer et al., 2009). Additionally, the inclusion of cellular variability in our simulations, achieved by incorporating the fluctuating dynamics of SNAIL, resulted in certain observable differences when compared to deterministic simulations: 1) individual cells had variable return times to an epithelial state, despite identical exposure dose and duration, and 2) cells were able to switch phenotype among E, M and hybrid E/M states, establishing a dynamic equilibrium in the phenotypic distribution.

Our analysis has many limitations. First, our mathematical model considered a reduced EMT regulatory network comprising of a few canonical EMT and MET drivers (SNAIL, ZEB, miR-200), although networks with teams of epithelial and mesenchymal genes have been identified (Hari et al., 2021). Second, our model of epigenetic regulation of miR-200 by ZEB is phenomenological and lacks the granularity of including different epigenetic enzymes recruited by one or more EMT/ MET-TFs, chromatin status of those EMT/MET-TFs (Chaffer et al., 2013) and the dynamics of heterochromatin. A few recent models have combined mechanistic dynamics of transcriptional and epigenetic control (Chen et al., 2021; Folguera-Blasco et al., 2019), building on previous attempts to explain epigenetic memory (Dodd et al., 2007). Third, by not considering cell division events, we have excluded the role of cell cycle in epigenetic regulation (Nashun et al., 2015). Fourth, we do not consider the impact of any extracellular changes during EMT such as increased matrix stiffness, which can give rise to mechanical memory due to mechanochemical feedback loops (Deng et al., 2021; Kumar et al., 2014; Price et al., 2021). Nonetheless, our model was able to – a) provide insight into how the dynamics of epigenetic changes during and post EMT induction can affect the reversibility to an epithelial cell state, corroborating with existing quantitative experimental and theoretical analysis; and b) explain experimentally observed timescale differences for EMT reversibility when cells were exposed to varying duration of TGFβ treatments of varying duration.

## Author contributions

MKJ (computational) and MT (experimental) formulated and supervised research. PJ (computational) and SC and KM (experimental) performed research. JTG and HL contributed to computational data analysis. All authors contributed to writing and editing of the manuscript.

## Code Availability

The codes used for simulation in this study can be accessed at: https://github.com/Paras-Jain20/EMT_Epigenetic_Decay

## Materials and Methods

### Cell Culture

MCF10A cells were a gift from Dr. Jeffrey Rosen and cultured as previously described (Toneff et al., 2016). Cells were treated with 5 ng/mL recombinant human TGFβ1 (R&D Systems, 240-B) prepared according to the manufacture’s specifications. Cells were passaged every three days during TGFβ1 treatment and after TGFβ1 withdrawal. TGFβ1 was added to fresh growth medium two days after passaging and upon passaging.

### RNA isolation and qPCR

For mRNA analyzed at days 0, 6 and 18, RNA was extracted from cells using TriZol (ThermoFisher, 15596026) and isolated using the RNeasy Mini Kit (Qiagen, 74104) with RNase-Free DNase (Qiagen, 79254). For mRNA analyzed at days 27, 36 and 45 and all miRNAs, RNA was extracted and isolated from cells using the miRNeasy Mini Kit (Qiagen, 217084) and treated with RNase-Free DNase (Qiagen, 79254).

Reverse transcription was performed on 500 ng total RNA using the High Capacity cDNA Reverse Transcription Kit with RNase Inhibitor (ThermoFisher, 4374966). Power Up SYBR Green Master Mix (ThermoFisher, A25776) was used to perform qPCR using 10 ng cDNA. All qPCR reactions were performed in triplicate. mRNA qPCR primers are included below (**Table 1**).

**Table 1:**
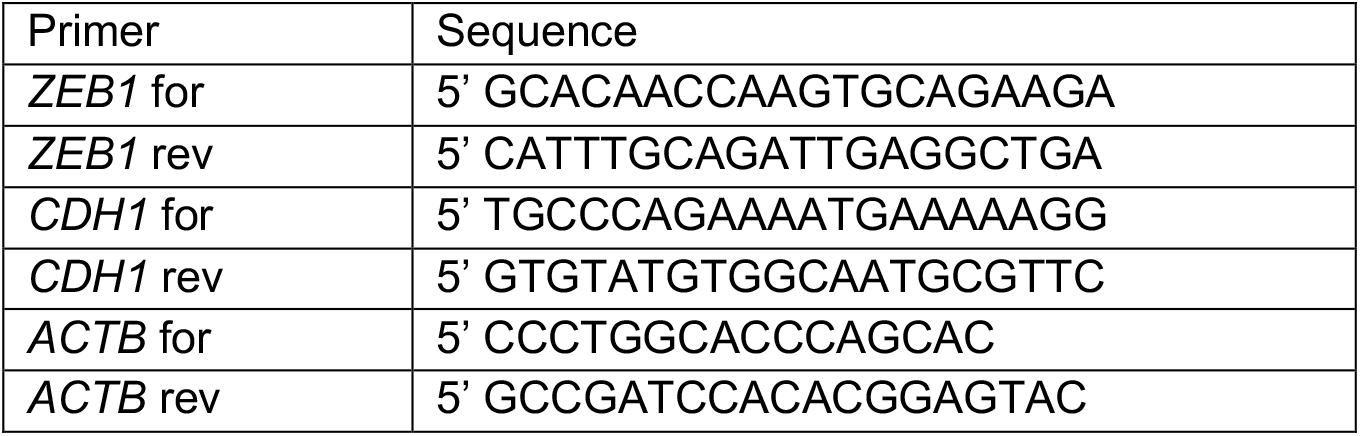
qPCR primer sequences.

Reverse transcription and qPCR for miRNAs was performed as previously described (Toneff et al., 2016). Taqman reverse transcription primers and qPCR probes (ThermoFisher) used were as follows: U6snRNA (001973), miR-200b (002251) and miR-200c (002300). Reverse transcription was performed using the Taqman microRNA Reverse Transcriptase Kit (ThermoFisher, 4366596). qPCR was performed using the TaqMan Fast Advanced Master Mix (ThermoFisher, 4444557).

All qPCR was performed using a CFX96 Touch Real-Time PCR System (BioRad).

### EMT regulatory network

We considered an EMT regulatory network involving interaction between canonical epithelial (miR-200) and mesenchymal (ZEB) markers with miR-200 and ZEB mutually repressing each other (**Fig 1A**, inset). SNAIL transcription factor acts an input to this network, supressing miR-200 and activating ZEB, and it represents cumulative effects of several EMT inducing signalling pathways, such as TGF-β, Wnt, and Notch (Lu et al., 2013). The rate equations capturing the production, degradation, and complex interactions between nodes for the network components are as follows:

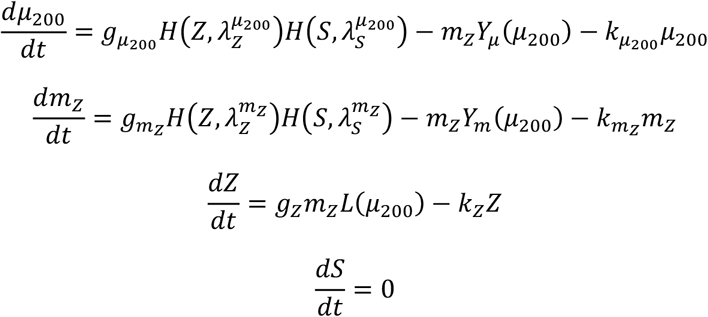

Here, *μ*_200_ = [miR-200], *m*_*z*_ = [ZEB1 mRNA], *Z* = [ZEB1], and *S* = [SNAI1]. [⋅] represents the concentration of a molecular species within a cell. *H* is the shifted Hill function.

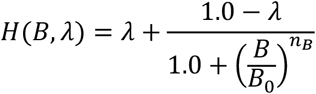

The functions *Y*_*μ*_, *Y*_*m*_, and *L* describe the post-transcriptional regulation of mRNA activity by micro-RNAs, as described earlier (Lu et al., 2013).

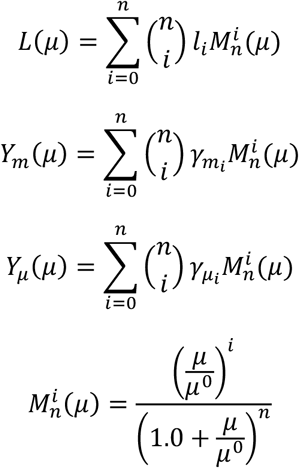

Here, *μ* is the microRNA concentration and *n* is number of micro-RNA binding sites on the mRNA. For the inhibition of *ZEB1* mRNA by miR-200, *n* = 6 and 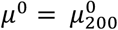. The values of all other kinetic parameters are listed in **Table 2** and **Table 3**.

**Table 2.** Regulatory network parameters – rates of production, and degradation; and Hills’ coefficient, threshold, and fold change for transcriptional regulations

| Parameter | Value | Parameter | Value |
| --- | --- | --- | --- |
| $g_{\mu_{200}}$ | $2.1 \times 10^3 \text{ mol. h}^{-1}$ | $n_Z^{\mu_{200}}$ | 3 |
| $g_{m_Z}$ | $11.0 \text{ mol. h}^{-1}$ | $n_S^{\mu_{200}}$ | 2 |
| $g_Z$ | $0.1 \times 10^3 \text{ h}^{-1}$ | $n_Z^{m_Z}$ | 2 |
| $k_{\mu_{200}}$ | $0.05 \text{ h}^{-1}$ | $n_S^{m_Z}$ | 2 |
| $k_{m_Z}$ | $0.5 \text{ h}^{-1}$ | $\lambda_Z^{\mu_{200}}$ | 0.1 |
| $k_Z$ | $0.1 \text{ h}^{-1}$ | $\lambda_S^{\mu_{200}}$ | 0.1 |
| $S_0^{\mu_{200}}$ | $180.0 \times 10^3 \text{ mol.}$ | $\lambda_Z^{m_Z}$ | 7.5 |
| $Z_0^{m_Z}$ | $25.0 \times 10^3 \text{ mol.}$ | $\lambda_S^{m_Z}$ | 10.0 |
| $S_0^{m_Z}$ | $180.0 \times 10^3 \text{ mol.}$ | $\mu_{200}^0$ | 10000 |
| $Z0u200^0$ | $220.0 \times 10^3 \text{ mol}$ | | |

**Table 3.** Parameters for mir200 – ZEB mRNA complex translation and degradation

| No. of miRNA binding sites | 0 | 1 | 2 | 3 | 4 | 5 | 6 |
| --- | --- | --- | --- | --- | --- | --- | --- |
| $l_i$ | 1.0 | 0.6 | 0.3 | 0.1 | 0.05 | 0.05 | 0.05 |
| $\gamma_{m_i} (\text{h}^{-1})$ | 0.0 | 0.04 | 0.2 | 1.0 | 1.0 | 1.0 | 1.0 |
| $\gamma_{\mu_i} (\text{h}^{-1})$ | 0.0 | 0.005 | 0.05 | 0.5 | 0.5 | 0.5 | 0.5 |

### Framework for incorporating epigenetic regulation of mir200 expression from ZEB

Epigenetic regulation of miR-200 can happen through increasingly methylation of its upstream promoter region, as noted during continued maintenance of mesenchymal state (Gregory et al., 2011). This epigenetic regulation can be incorporated phenomenologically by strengthening the miR-200 suppression by ZEB depending on the duration for which ZEB levels are maintained high. To model this process, we earlier modelled the threshold of repression of miR-200 by ZEB a dynamic variable whose rate of change depended on ZEB levels (Jia et al., 2019), as shown below:

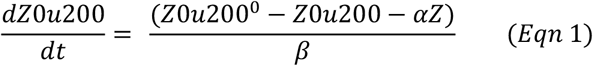

Where,

*Z*0*u*200 : threshold for transcriptional repression

*Z*0*u*200^0^ : basal threshold (constant parameter)

*Z* : ZEB levels

*β*: time constant (constant parameter)

*α* : epigenetic regulation strength (constant parameter)

In simulations shown here, the basal threshold *Z*0*u*200^0^ = 220 × 10^3^ molecules (Lu et al., 2013), and the epigenetic regulation strength parameter, *α* = 0.15, corresponding to a strong epigenetic regulation. *α* values greater than 0.15 gives a negative value for Z0u200 levels within relevant range of SNAIL levels (0-600 × 10^3^ molecules), thus becoming biologically inappropriate. The time constant *β* scales the response time of threshold *Z*0*u*200 to changes in ZEB levels. To take into account any possible differences in the molecular mechanisms and/or reaction rates of epigenetic changes during EMT and its reversal, we consider two independent *β* values – 1) *β*_*for*_, during induction, and 2) *β*_*rev*_, during withdrawal. To analyse the cellular response without epigenetic regulation, we set *α* = 0, *β*_*for*,_ = 1 *hr*, and *β*_*rev*_ = 1 *hr*.

### EMT induction and withdrawal simulation set-up

SNAIL levels are being used to control the induction and reversal of EMT, based on bifurcation diagram (**Fig 2A**). These levels can be affected by its stochastic gene expression, variability in upstream signalling pathway activity, and varying microenvironmental cues (Strasen et al., 2018). To model the temporal variability of SNAIL levels, we represented its dynamics using Ornstein-Uhlenbeck (OU) process, as mentioned earlier (Li et al., 2016):

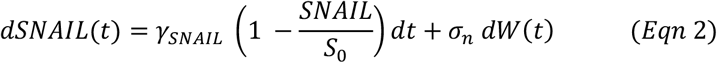

Here, *γ*_SNAIL_: return rate, *S*_0_: mean cellular SNAIL level, W(*t*) : Weiner process, *σ*_*n*_: standard deviation of noise. With the above equation, SNAIL levels follow a stochastic trajectory whose statistical characteristics at stationary state are highlighted in **Table 4**.

**Table 4.**
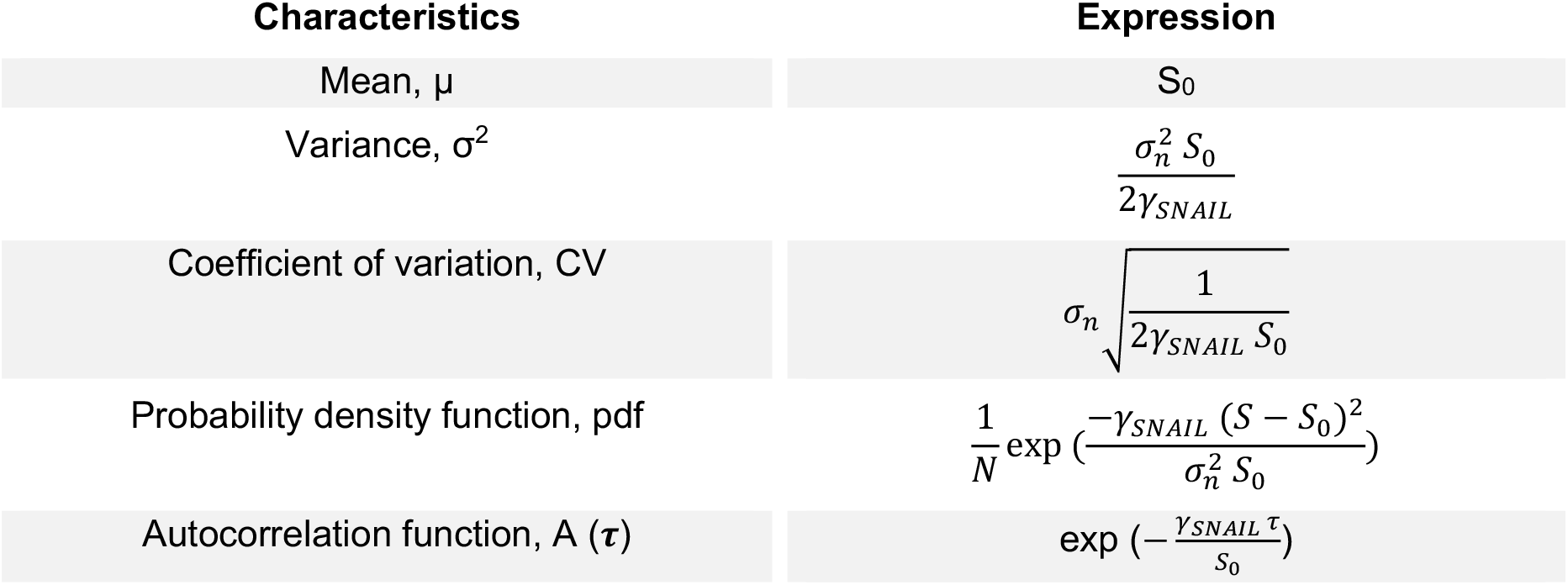
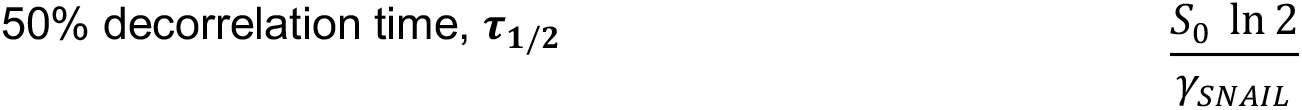
Statistical characteristic of stochastic SNAIL dynamics (at stationary state)

In the simulations:

1. For deterministic SNAIL dynamics (**Fig 2-5**), *σ*_*n*_ = 0.
2. *S*_0_ attains two values:
  a. *S*_01_ (pre-induction/post-withdrawal levels): mean SNAIL level of a cell in the population prior to EMT induction and to which it returns after a prolonged withdrawal period.
  b. *S*_02_ (post-induction levels): mean SNAIL level of a cell in the population after long duration of EMT induction.
3. *τ*_1/2_ = 120 hrs (5 days) following experimental studies which report mixing times of proteins in human cancer cell lines (Li et al., 2016; Sigal et al., 2006). We considered *τ*_1/2_ to remain conserved irrespective of the mean SNAIL level of a cell (S_0_).
4. With the above constraint, the return rate (*γ*_*SNAIL*_) in SNAIL’s dynamics gets defined:

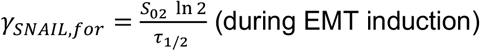

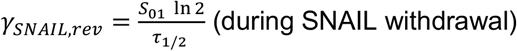
5. For stochastic (noisy) SNAIL dynamics, the choice of standard deviation of noise in SNAIL dynamics, *σ*_*n*_, is made by considering the relation between CV, *τ*_1/2_, S_0_, and *σ*_*n*_ values derived by eliminating *α*_SNAIL_ from CV and *τ*_1/2_ expressions (**Table 4**) as shown below:

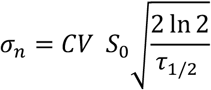 Note that the CV of SNAIL levels is inversely proportional to square root of S_0_ (Table 3). Here, CV value is taken as 0.3 at S_0_ = 150,000 molecules. This choice of CV lies within the biological observed range for CVs of several proteins from variety of pathways (Sigal et al., 2006). Substituting numerical values of CV = 0.3, S_0_ = 150,000 molecules, and τ_1/2_ = 120 *hr* above, we get:

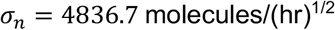 The above value *σ*_*n*_ is used during both EMT induction and withdrawal time points for all stochastic population-level simulations.

For population-level analysis (Fig. 6 and corresponding SI figures), we simultaneously generate 10000 independent trajectories of SNAIL (Eqn. 2), each representing an individual cell (**Fig. 6A**).

#### Simulation procedure

A cell in the model is described by 6 variables: {miR-200, ZEB mRNA, ZEB, SNAIL, Z0u200 and S_0_}. The ZEB mRNA levels, as per bifurcation diagram, are used to assign phenotypes (**Table 5**).

**Table 5.**
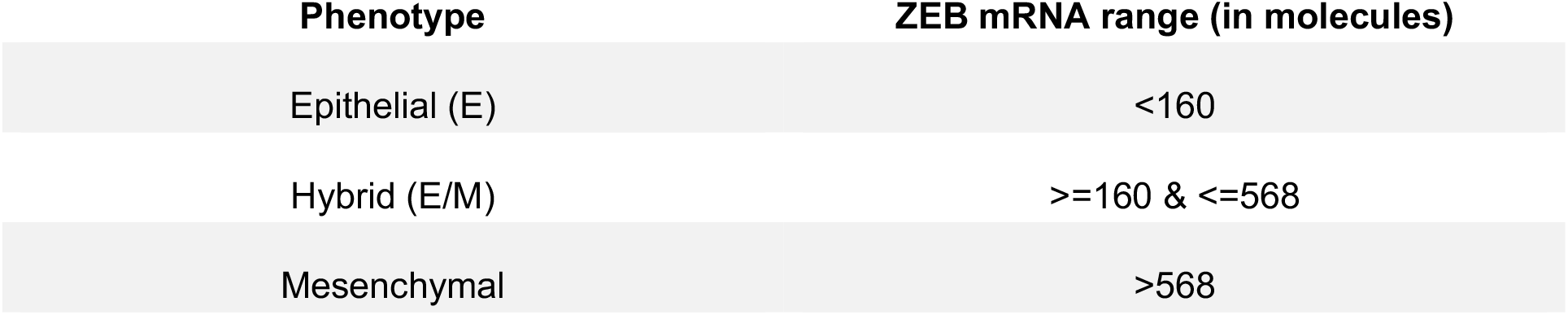
ZEB mRNA ranges and cell’s phenotype

#### For deterministic single-cell EMT induction and withdrawal

1. Initialize model parameters: Assign values to *α*, τ_1/2_, *β*_*for*,_, *β*_*rev*_, *γ*_*SNAIL,for*,_, *γ*_*SNAIL,rev*_, S_01_, and S_02_ either using the user inputs or relations described in section 3. Set σ_2_ = 0.
2. Initialize cellular network components: Set SNAIL = S_01_, Z0u200 = Z0u200^0^, mir200 = 0, ZEB mRNA = 0, and ZEB = 0. Simulate the model for an arbitrary long time so that variables settle to their steady values for the given SNAIL level.
3. Define the time for EMT induction and SNAIL withdrawal.
4. For EMT induction, assign S_0_ = S_02_, *β* = *β*_*for*,_, and *γ*_*SNAIL*_ = *γ*_*SNAIL,for*,_. Then simulate the model for defined number of induction days.
5. When induction duration ends, assign S_0_ = S_01_, *β* = *β*_*rev*_, and *γ*_*SNAIL*_ = *γ*_*SNAIL,rev*_. Then simulate the model for defined number of withdrawal days.

#### For stochastic population-level EMT induction and withdrawal

1. Initialize model parameters: Assign values to *α, CV, S*_0_*CV*^**#**^, τ_1/2_, *β*_*for*,_, *β*_*rev*_, *γ*_*SNAIL,for*,_, *γ*_SNAIL,*rev*_, *S*_*01*_, *S*_*02*_, and *σ*_*n*_ either using the user inputs or relations described in section 3.
2. Generate 10K normally distributed SNAIL samples centred around S_0_ = S_01_ with variance, 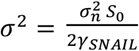. Assign one SNAIL sample to each cell with the values of the other cellular network components as mir200 = 0, ZEB mRNA = 0, ZEB = 0, Z0u200 = Z0u200^0^, and S_0_ = sampled SNAIL value. Simulate the model for an 1000 hours with *σ*_*n*_ = 0 so that variables settle to their steady values for the given SNAIL levels.
3. Again, for 1000 hours, run the system while considering noise in SNAIL dynamics (use *σ*_*n*_ as determined above).
4. Define the time for EMT induction and SNAIL withdrawal.
5. For EMT induction, assign S_0_ = S_02_ for every cell, *β* = *β*_*for*,_, and *γ*_*SNAIL*_ = *γ*_*SNAIL,for*,_. Then simulate the model for defined number of induction days.
6. When induction duration ends, assign S_0_ = S_01_ for every cell, *β* = *β*_*rev*_, and *γ*_SNAIL_ = *γ*_SNAIL,*rev*_. Then simulate the model for defined number of withdrawal days.

^**#**^*S*_0_*CV* : *S*_0_ value for which CV value is initialized

Here, *mol*. ≡ *molecules* / *cell*

Where, *N* is a normalising factor, and τ is lag duration. τ_1/2_ represents the average time at which a cell’s SNAIL level changes by 50% due to stochasticity in its expression.

## Supplementary figures

**Fig S1.**
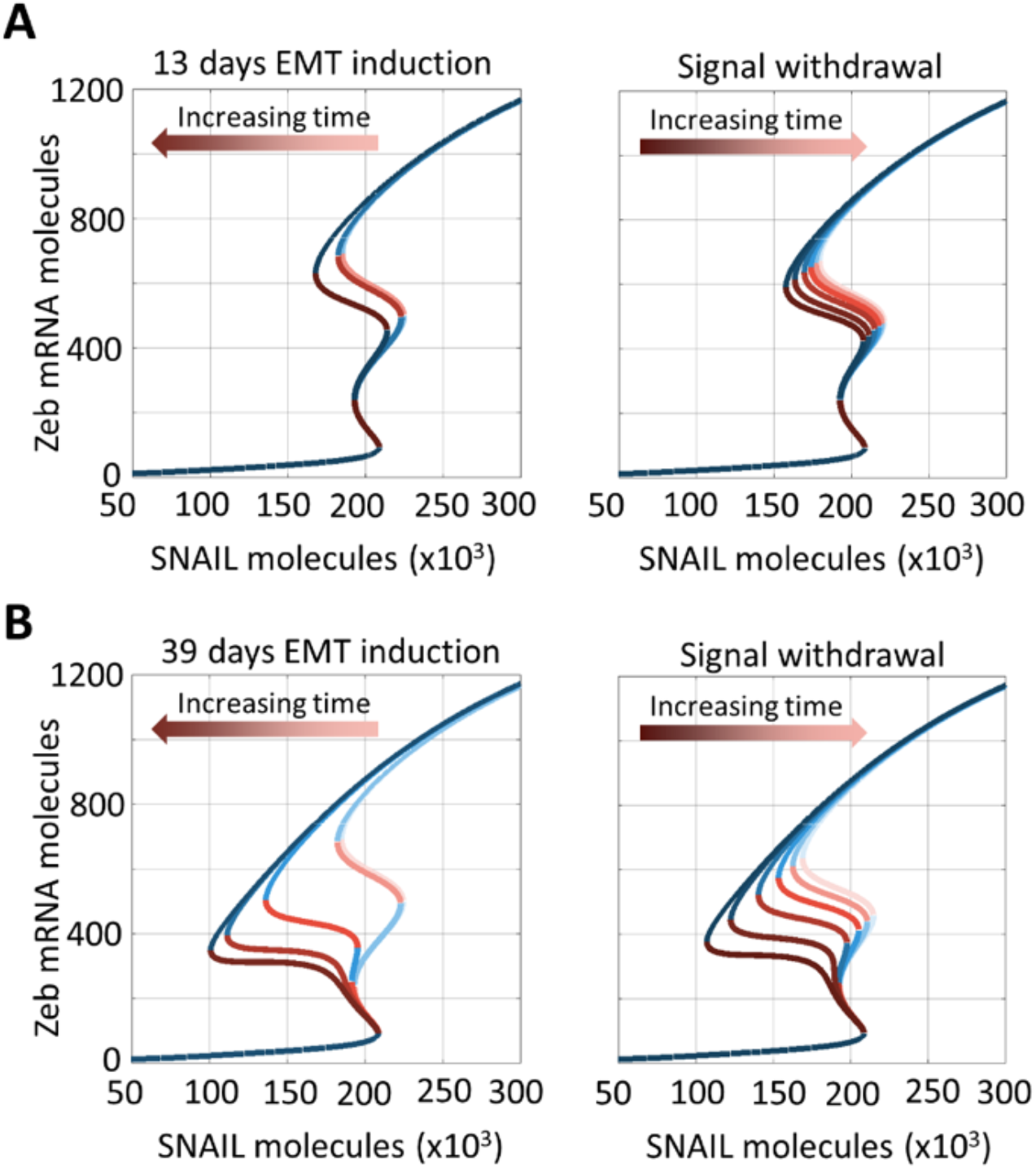
Effect of epigenetic regulation on phenotypic stability as a function of EMT induction period. Bifurcation diagrams for ZEB mRNA levels for varying strengths of epigenetic regulation (Z0u200 levels) during EMT induction (left) or withdrawal (right) of EMT-inducing signal at **A**) (from left to right) 0, 10, 13 days post-induction and (from right to left) 10, 20, 30, 40, 50, 60 days post-withdrawal, **B**) (from left to right) 0, 10, 20, 30, 39 days post-induction and (from right to left) 10, 20, 30, 40, 50, 60 days post-withdrawal.

**Fig S2.**
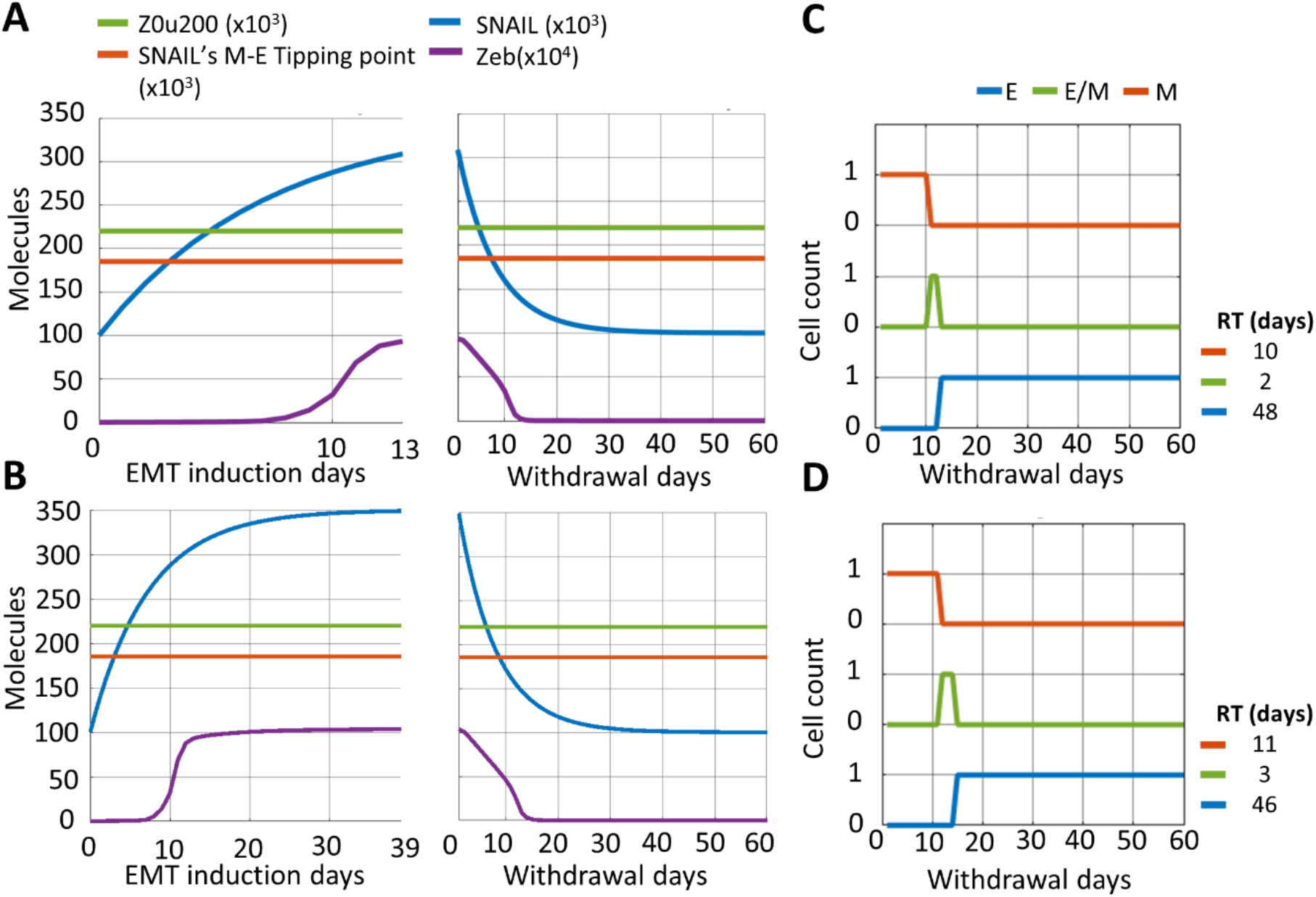
Influence of duration of EMT induction on time to recover an epithelial state post-withdrawal without considering any epigenetic regulation. Dynamics of ZEB levels and changes in epigenetic regulation measures – Z0u200 and SNAIL’s M-E tipping levels – during EMT induction for **A)** 13 days, **B)** 39 days followed by withdrawal period of 60 days. **C** and **D** show change in cell’s phenotype during withdrawal period after EMT induction for 13 days (A) and 39 days (B), respectively. Parameters used to obtain results from A-D are: α = 0, S_01_ = 100K molecules, S_02_ = 350K molecules.

**Fig S3.**
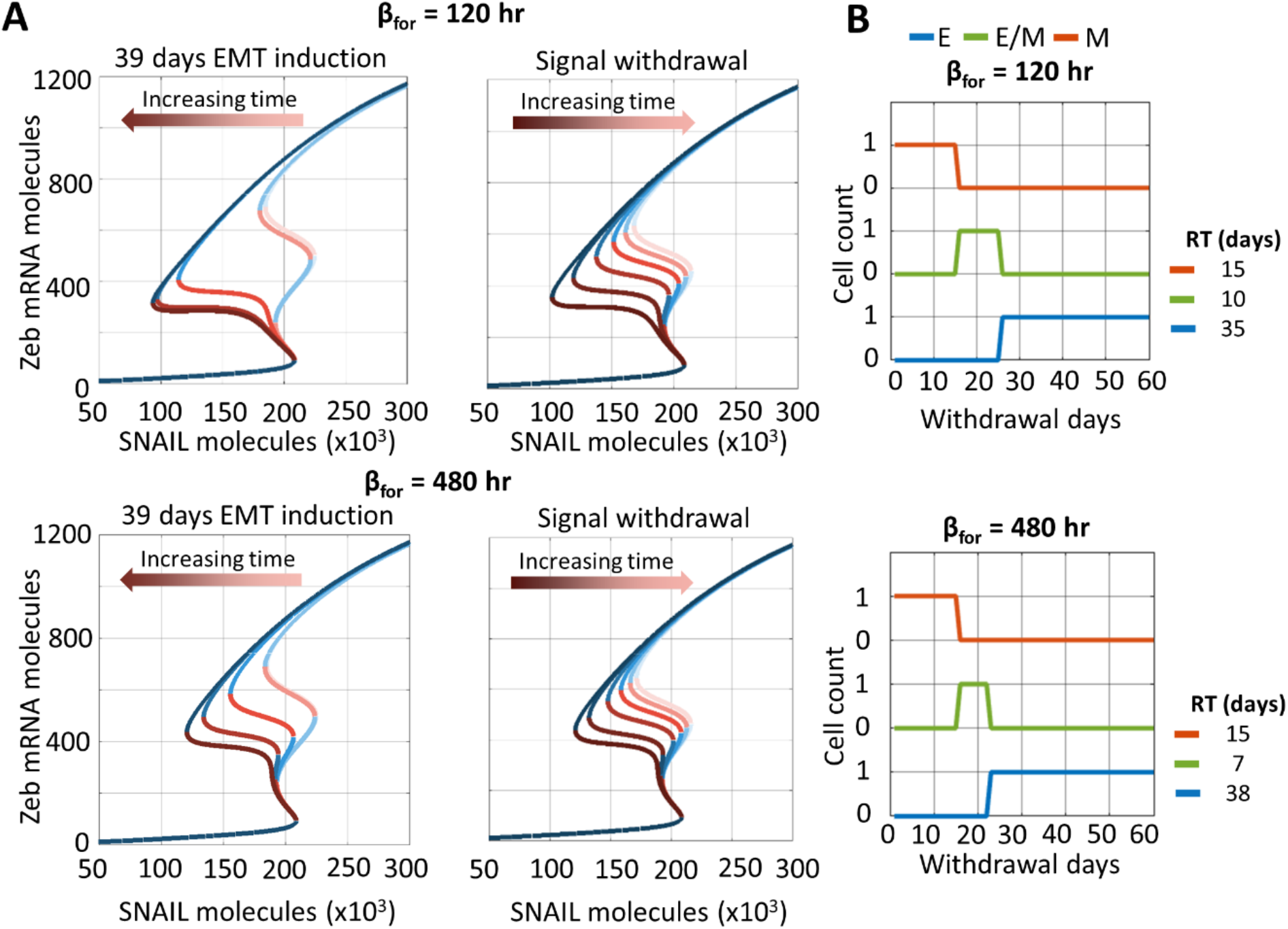
Effect of varying the rates of accumulation of epigenetic memory during EMT on the phenotypic stability and time to recover an epithelial state. **A)** Bifurcation diagrams for ZEB mRNA levels for temporally varying effect of epigenetic regulation (decreasing or increasing Z0u200 levels) during EMT induction (left) or withdrawal (right) of EMT-inducing signal at 0, 10, 20, 30, 39 days post-induction and 10, 20, 30, 40, 50, 60 days post-withdrawal when β_for_ = 120 and 480 hr. **B)** Cell state switching post EMT-inducer withdrawal.

**Fig S4.**
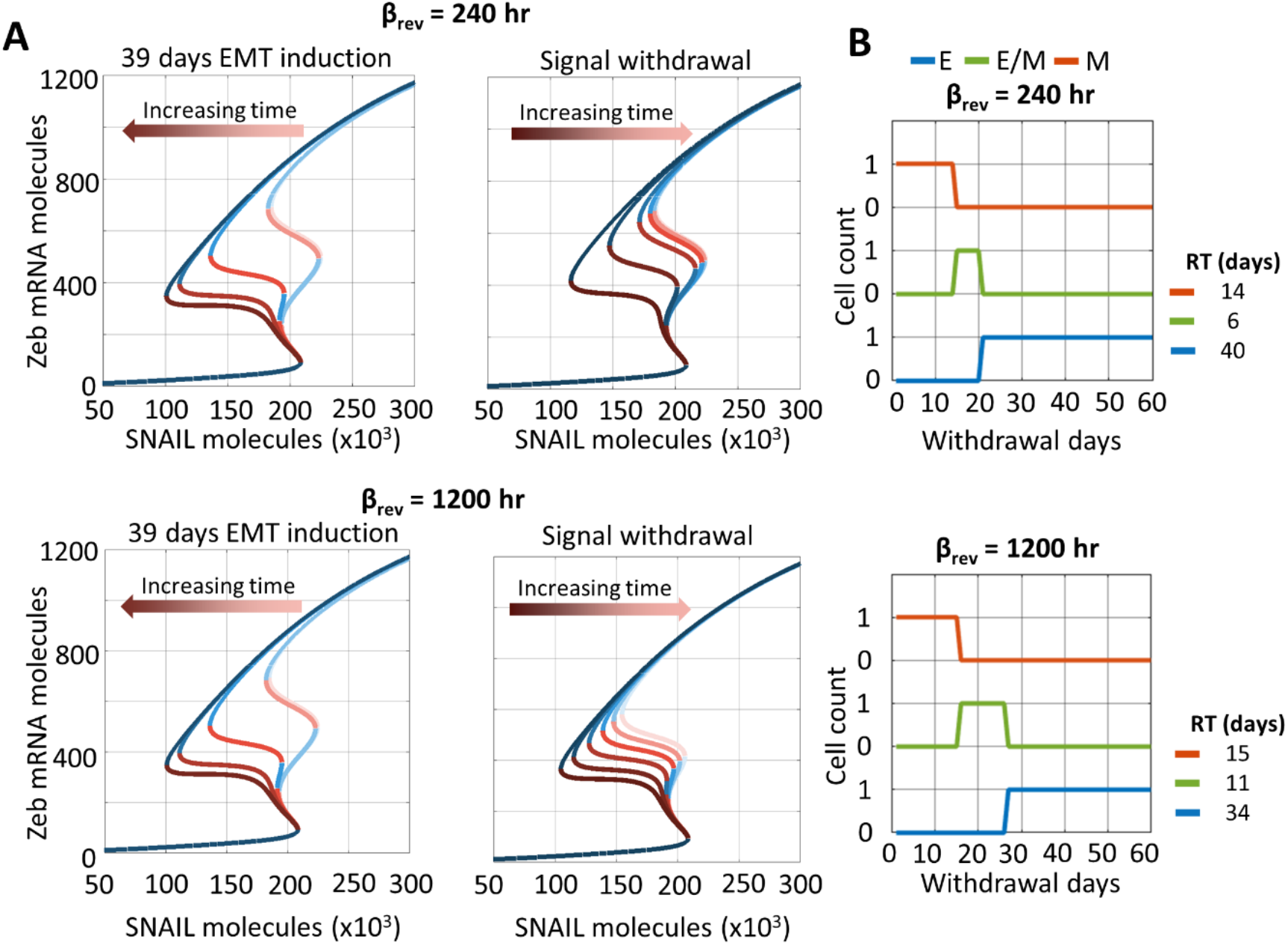
Effect of varying rates of decay of epigenetic memory during withdrawal on phenotypic stability and time to recovery epithelial state. **A)** Bifurcation diagrams for ZEB mRNA levels for temporally varying effect of epigenetic regulation during EMT induction (left) or withdrawal (right) of EMT-inducing signal at 0, 10, 20, 30, 39 days post-induction and 10, 20, 30, 40, 50, 60 days post-withdrawal when β_rev_ = 240 and 1200 hr. **B)** Cell state switching post EMT-inducer withdrawal.

**Fig S5.**
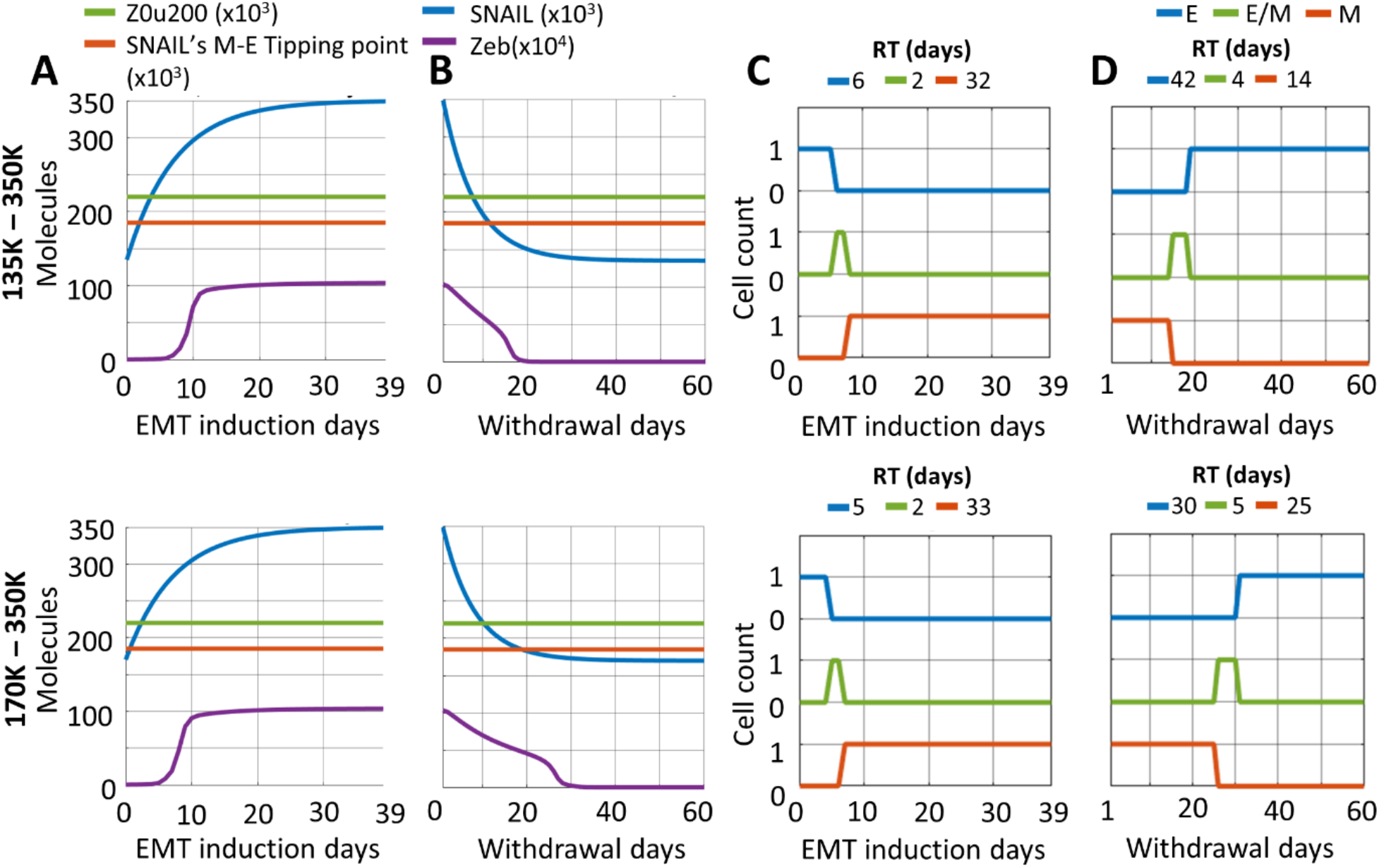
Return times to epithelial state with variation in pre-induction (post-withdrawal) SNAIL levels (S_01_), without considering epigenetic changes during EMT. **A, B)** Changes in SNAIL, and ZEB during EMT and withdrawal period. **C, D**) Change in cell’s phenotype during EMT and withdrawal period. The results were generated considering the parameters: α = 0 (no epigenetic regulation), S_02_ = 350K molecules, and varying S_01_.

**Fig S6.**
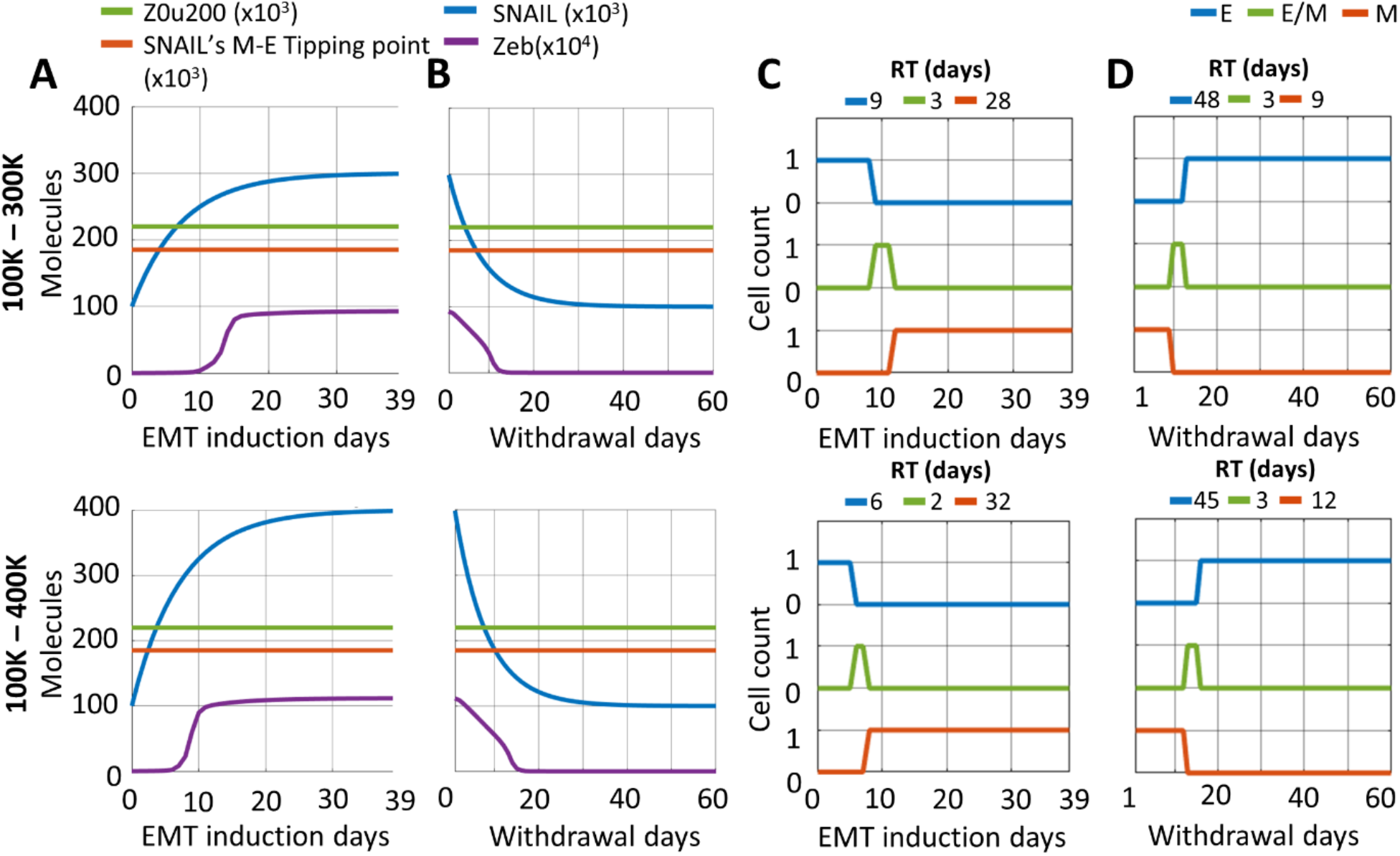
Return times to epithelial state with variation in post-induction (S_02_) SNAIL levels, without considering epigenetic changes during EMT. **A, B)** Changes in SNAIL, and ZEB during EMT and withdrawal period. **C, D**) Change in cell’s phenotype during EMT and withdrawal period. The results were generated considering the parameters: α = 0 (no epigenetic regulation), S_01_ = 100K molecules, and varying S_02_.

**Fig S7.**
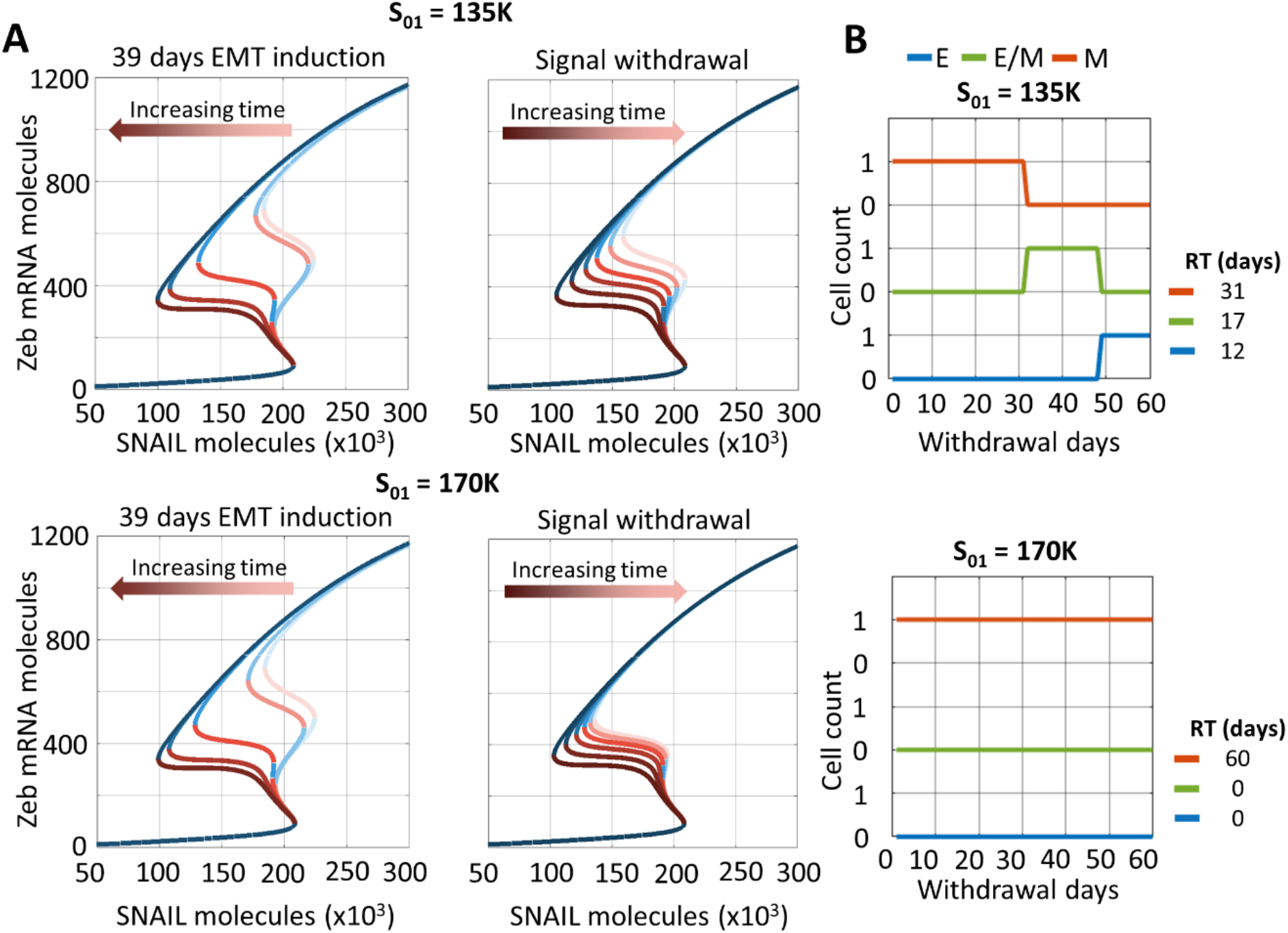
Effects of varying pre-induction (post-withdrawal) SNAIL levels (S_01_) on phenotypic stability and time to recovery epithelial state in the presence of epigenetic regulation. **A)** Bifurcation diagrams for ZEB mRNA levels for temporally varying effects of epigenetic regulation (decreasing or increasing Z0u200 levels) during EMT induction (left) or withdrawal (right) of EMT-inducing signal at 0, 10, 20, 30, 39 days post-induction and 10, 20, 30, 40, 50, 60 days post-withdrawal when S_01_ = 135K and 170K molecules. **B)** Cell state switching post EMT-inducer withdrawal.

**Fig S8.**
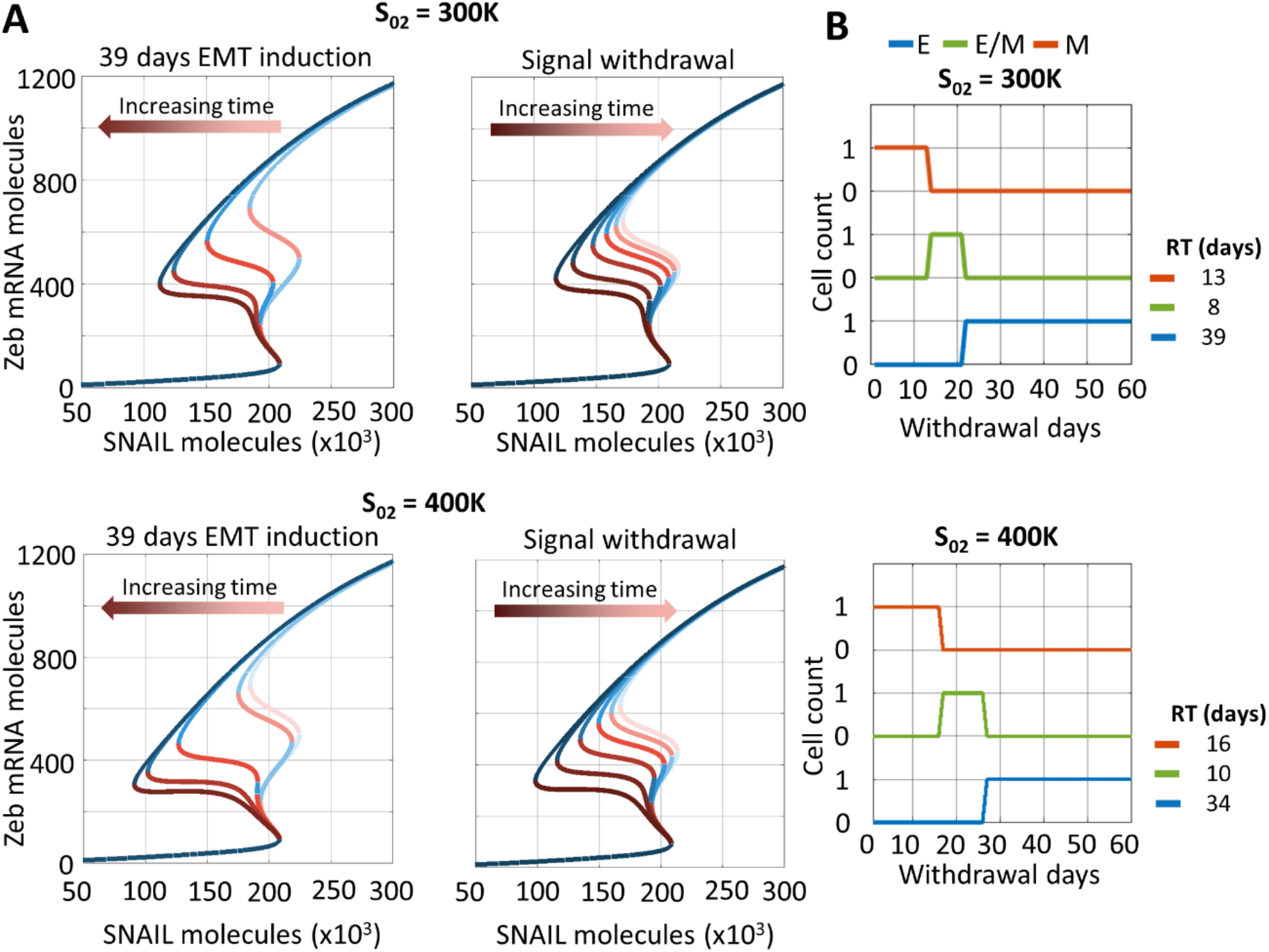
Effects of varying post-induction SNAIL levels (S_02_) on phenotypic stability and time to recovery epithelial state in presence of epigenetic regulation. **A)** Bifurcation diagrams for ZEB mRNA levels for temporally varying effects of epigenetic regulation (decreasing or increasing Z0u200 levels) during EMT induction (left) or withdrawal (right) of EMT-inducing signal at 0, 10, 20, 30, 39 days post-induction and 10, 20, 30, 40, 50, 60 days post-withdrawal when S_02_ = 300K and 400K molecules. **B)** Cell state switching post EMT-inducer withdrawal.

**Fig S9.**
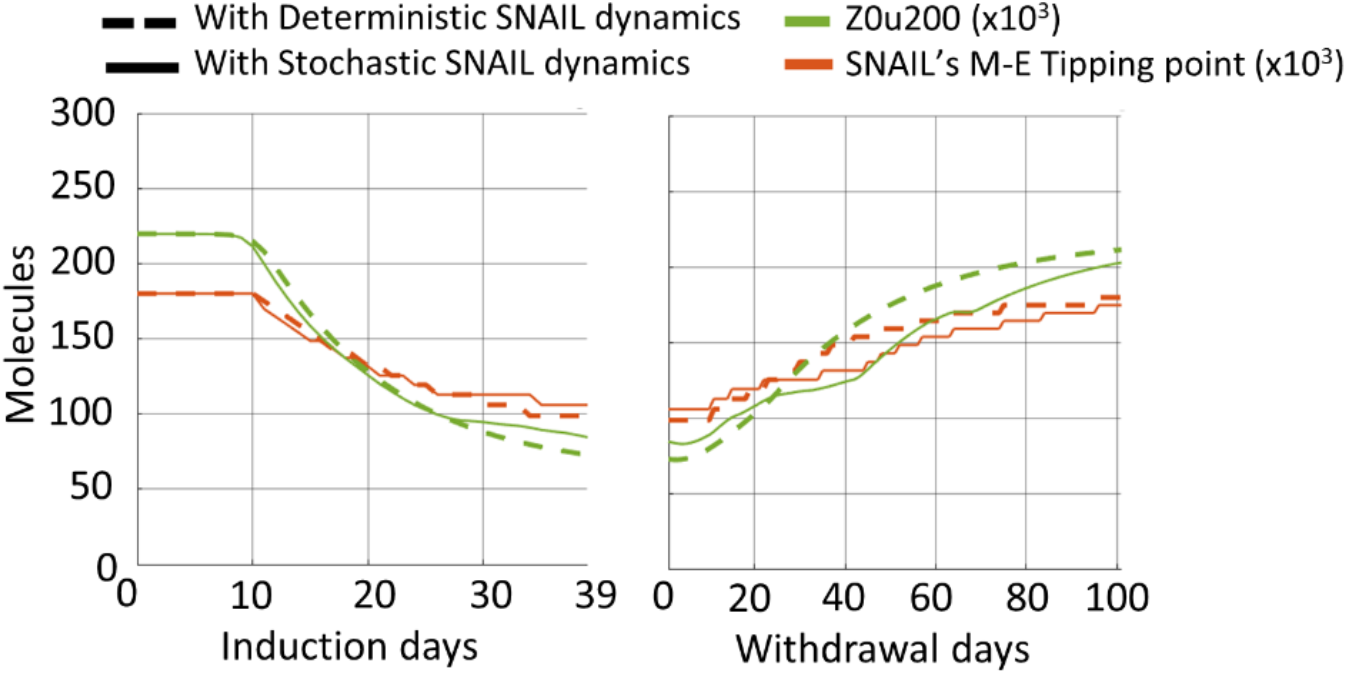
Effects of stochastic SNAIL dynamics on epigenetic regulation measures –Z0u200 and SNAIL’s M-E tipping level – during EMT and withdrawal. Changes in Z0u200 and SNAIL’s M-E tipping level during EMT induction and withdrawal for stochastic (solid curves) and deterministic (dashed curves) dynamics of cellular SNAIL levels (Fig. 6D). Plots generated using following parameters for simulations: α = 0.15, S_01_ = 100K molecules, S_02_ = 350K molecules, β_for_ = 240 hr, and β_rev_ = 720 hr.

**Fig S10.**
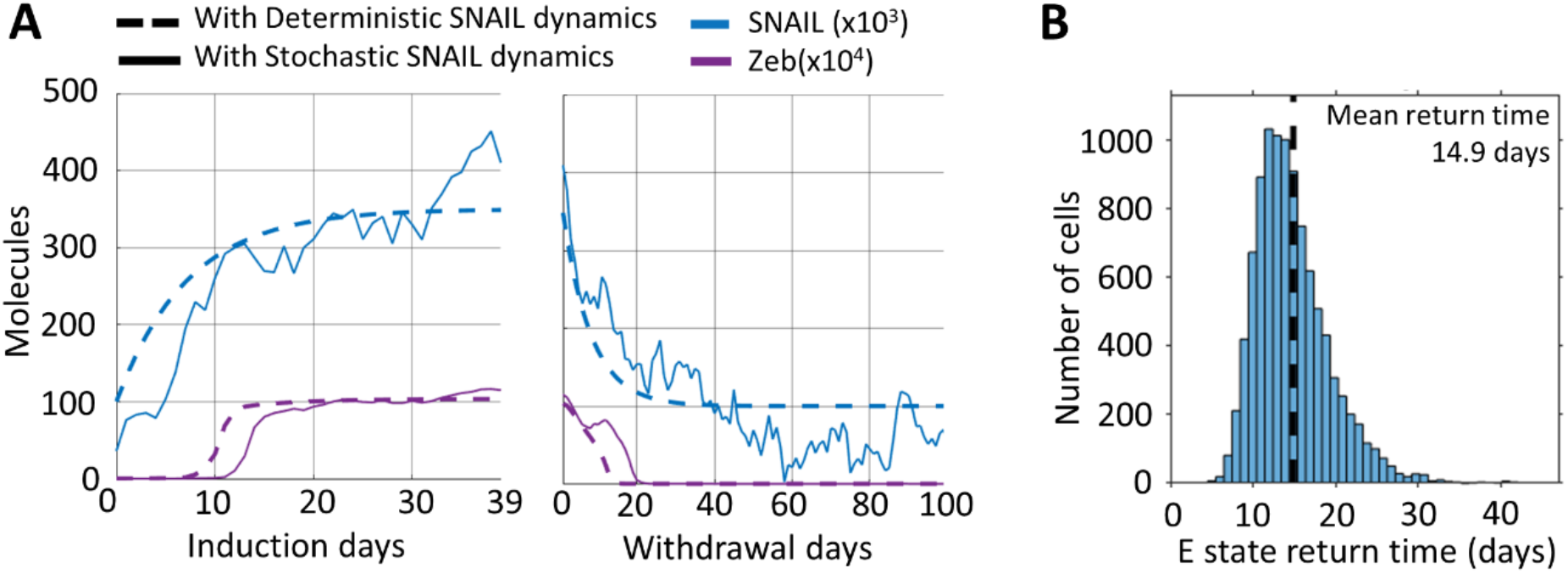
Effects of stochastic SNAIL dynamics on cellular ZEB levels, and epithelial state recovery time of individual cells in the population, without considering epigenetic changes during EMT. **A)** Changes in SNAIL and ZEB levels during EMT induction and withdrawal period for a cell in the population, with and without stochastic fluctuations in SNAIL expression, shown by solid and dashed curves, respectively. Deviation in ZEB levels were observed on comparing stochastic (noisy) and deterministic (noise-less) SNAIL dynamics. **B)** Distribution of return time of cells to the epithelial state after undergoing EMT induction for 39 days. Population size was 10K cells, and results were obtained using parameters α = 0, S_01_ = 100K molecules, and S_02_ = 350K molecules, which were assigned to every cell in the population.

**Fig S11.**
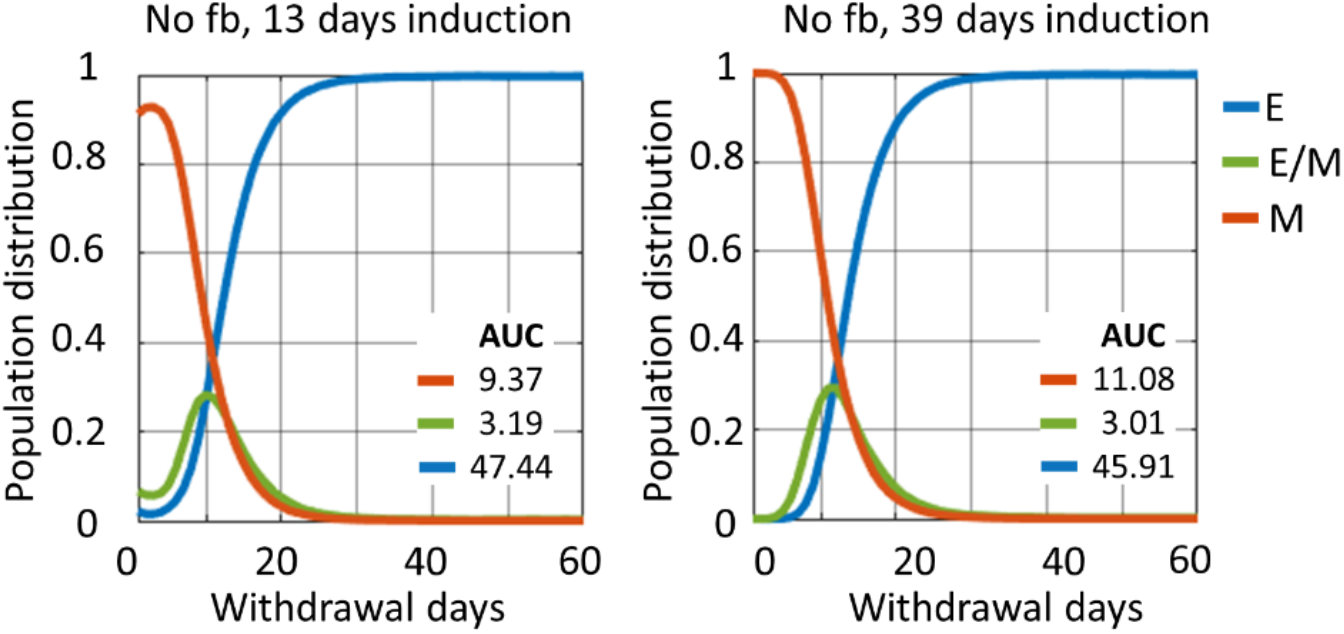
Change in phenotypic distribution of cell population during withdrawal period after 13 and 39 days of EMT induction, while epigenetic changes during EMT weren’t not considered. We observed enhanced residence time of cells in M states during withdrawal for 39 days when compared to 13 days EMT induction (increasing AUC of M state). The results presented were mean of three independent simulation runs of 10K cells each. Plots generated using following parameters for simulations: α = 0, S_01_ = 100K molecules, and S_02_ = 350K molecules.

**Fig S12.**
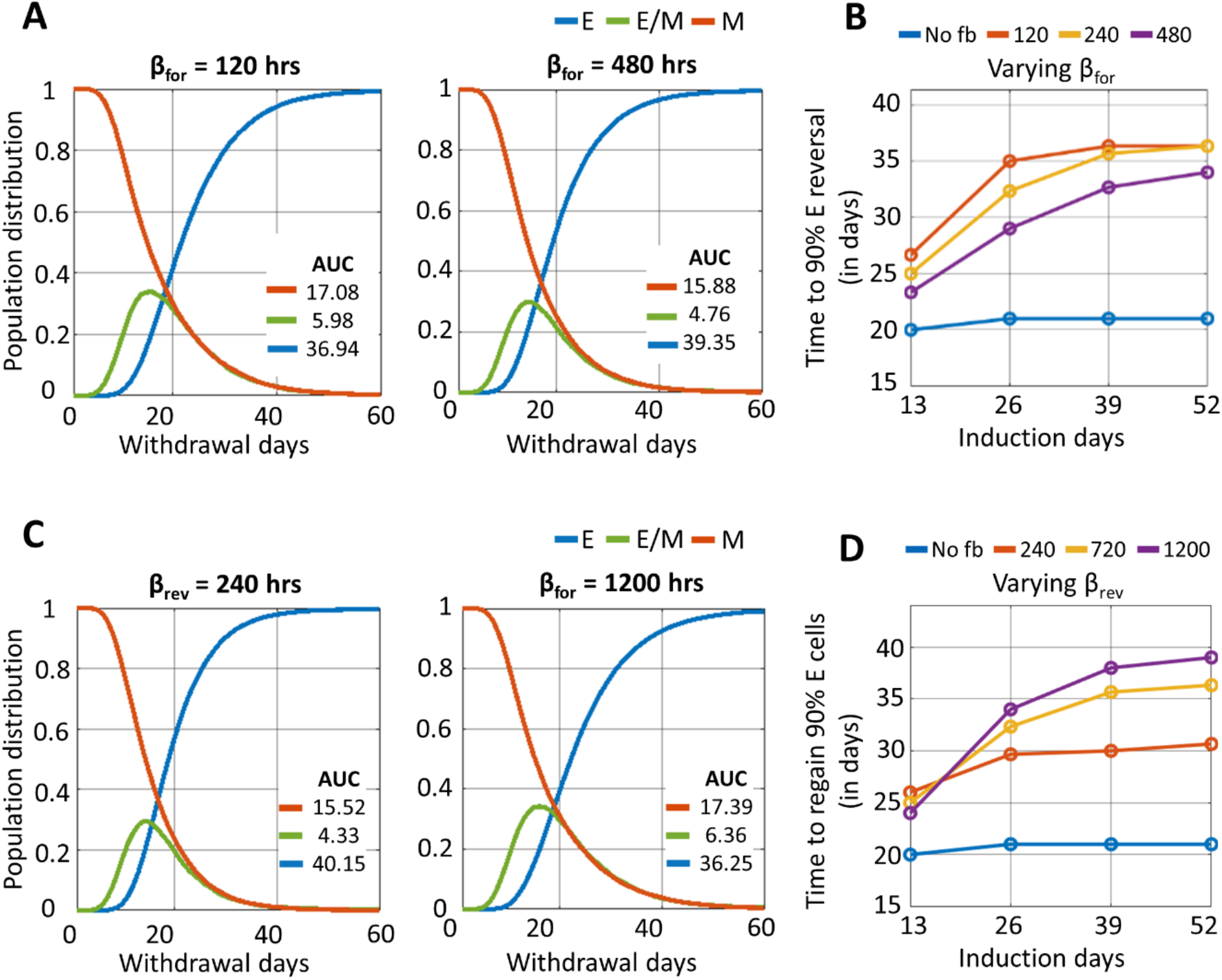
Effects of varying epigenetic memory accumulation and decay rates on epithelial recovery dynamics in a cell population. **A)** Change in phenotypic distribution of cell population during withdrawal period after 39 days of EMT induction with varying β_for_ values. Lower β_for_ caused faster epigenetic changes during EMT, and therefore, enhanced the residence time of cells in M and E/M states (increasing AUC of M and E/M states with decreasing β_for_ values). **B)** Time taken to regain 90% of epithelial phenotype share in the population on varying both the durations of EMT induction and β_for_. **C)** Change in phenotypic distribution of cell population during withdrawal period after 39 days of EMT induction with varying β_rev_ values. Higher β_rev_ cause slower decay of epigenetic changes acquired during EMT, and therefore, enhance the residence time of cells in M and E/M states (increasing AUC of M and E/M states with higher β_rev_ values). **D)** Time taken to regain 90% of epithelial phenotype share in the population on varying both the durations of EMT induction and β_rev_. The results presented were mean of three independent simulation runs of 10K cells each. Base parameters used for simulations were α = 0.15, S_01_ = 100K molecules, and S_02_ = 350K molecules, β_for_ = 240 hrs, and β_rev_ = 720 hrs, with specific ones being perturbed in their corresponding cases as is shown in the figure panel’s title or legend (A-D). The above parameters were assigned to every cell in the population.

**Fig S13.**
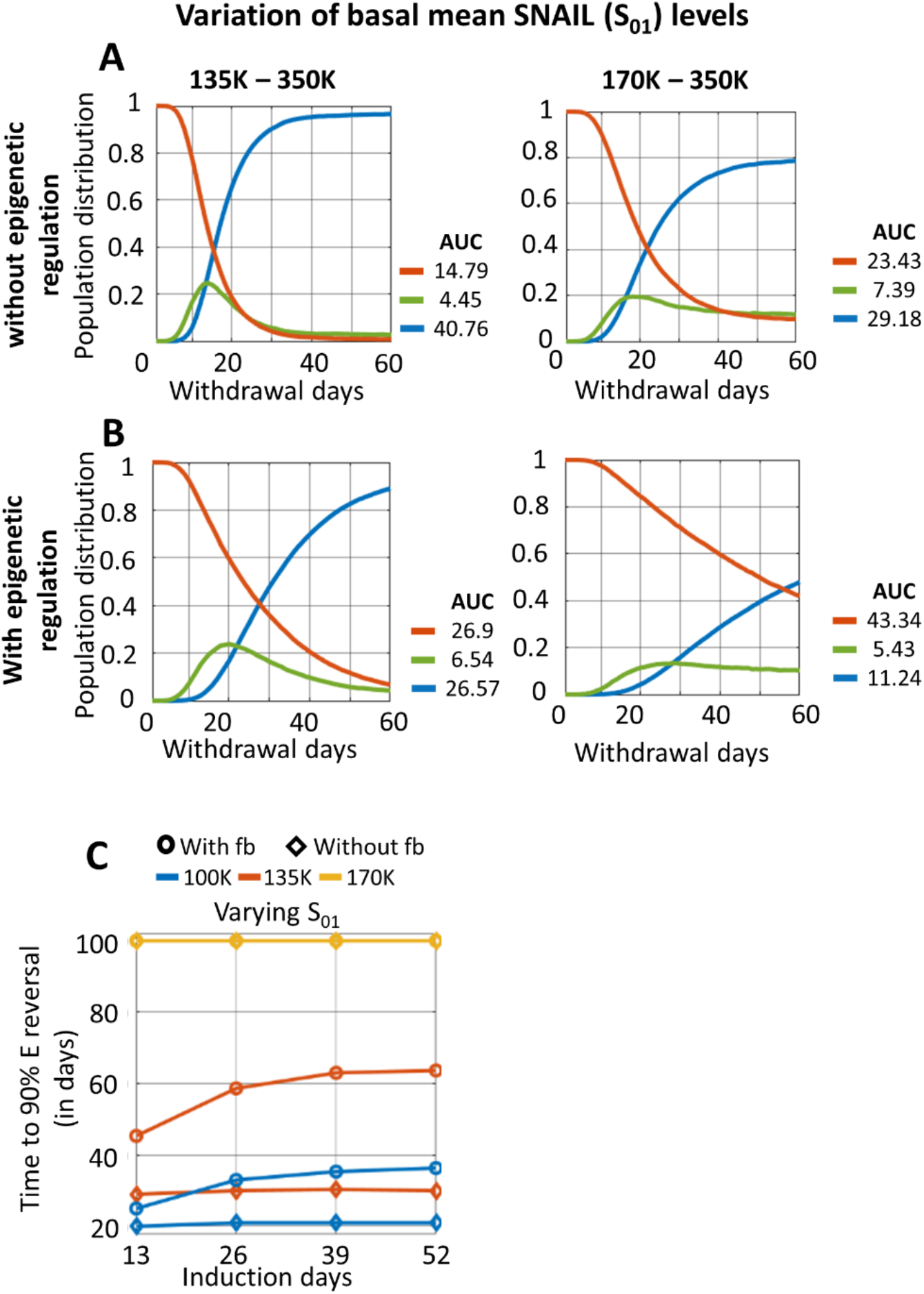
Effects of varying mean pre-induction (post-withdrawal) SNAIL levels (S_01_) of the population on epithelial state recovery dynamics in a cell population. **A)** Change in phenotypic distribution of cell population during withdrawal period after 39 days of EMT induction with varying S_01_, while epigenetic changes during EMT weren’t considered. Increasing S_01_ slowed down E fraction recovery and changed the steady state phenotypic distribution of the cell population. **B)** Change in phenotypic distribution of cell population during withdrawal period after 39 days of EMT induction with varying S_01_, while epigenetic changes during EMT being considered. With epigenetic regulation, higher S_01_ further slowed down epithelial state recovery dynamics. **C)** Time taken to regain 90% of epithelial phenotype share in the population on varying both the durations of EMT induction and S_01_. The results are presented for both with and without epigenetic regulation scenarios. Increasing period of EMT delayed epithelial state recovery for a given S_01_ values when epigenetic changes occur during EMT. For S_01_ = 170K, the 90% recovery time was >100 days for all durations of EMT induction, as the steady state percent of epithelial state was below 90% (end time point of plot B). The results presented were mean of three independent simulation runs of 10K cells each. Base parameters used for simulations were α = 0.15, S_01_ = 100K molecules, and S_02_ = 350K molecules, β_for_ = 240 hrs, and β_rev_ = 720 hrs, with specific ones being perturbed in their corresponding cases as is shown in the figure panel’s title or legend (A-E). The above parameters were assigned to every cell in the population. For cases without epigenetic regulation, α = 0.

**Fig S14.**
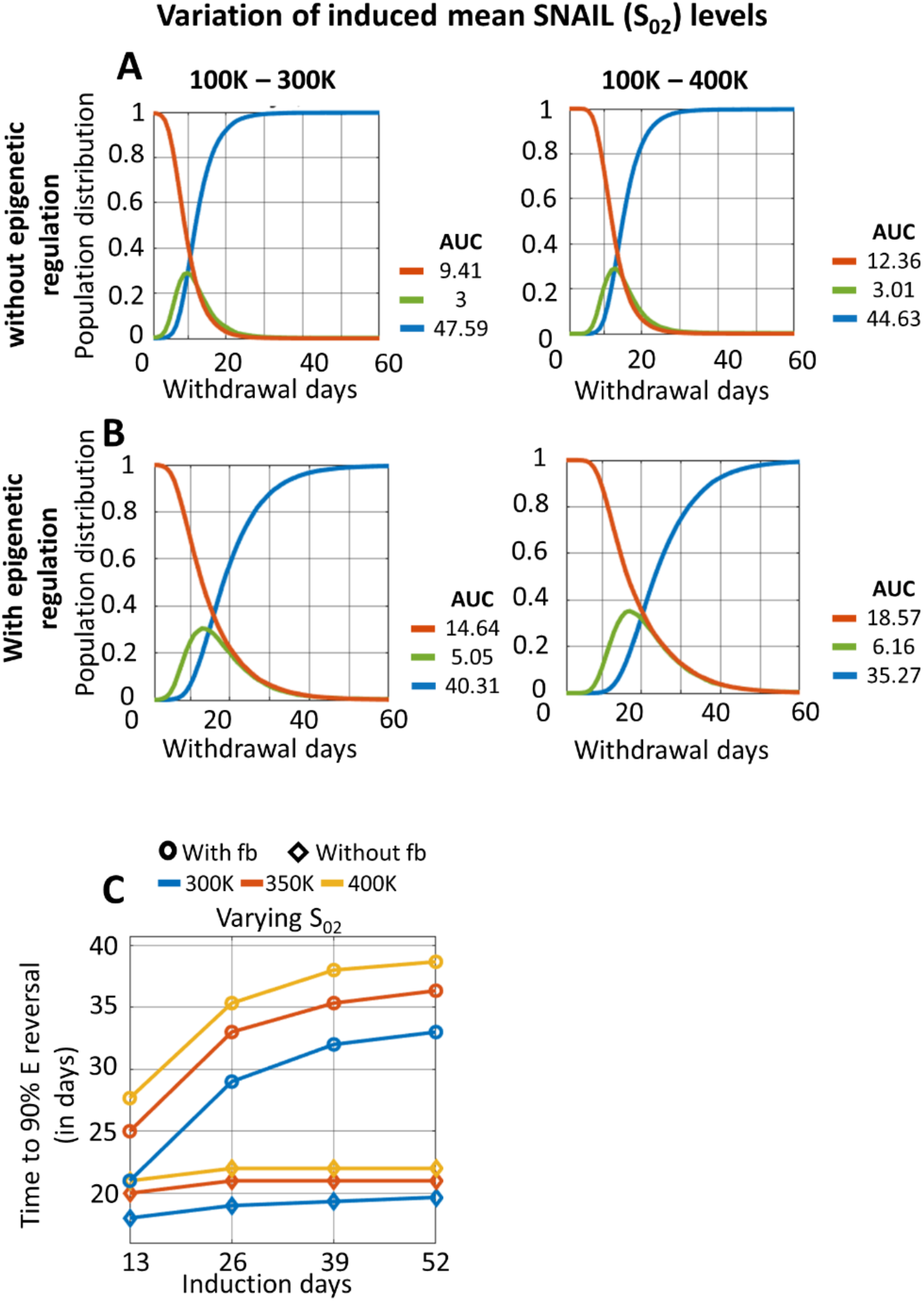
Effects of varying mean post-induction SNAIL levels (S_02_) of the population one epithelial state recovery dynamics in a cell population. **A)** Change in phenotypic distribution of cell population during withdrawal period after 39 days of EMT induction with varying S_02_, while epigenetic changes during EMT weren’t considered. Increasing S_02_ slowed down E fraction recovery. **B)** Change in phenotypic distribution of cell population during withdrawal period after 39 days of EMT induction with varying S_02_, while epigenetic changes during EMT being considered. With epigenetic regulation, higher S_02_ further slowed down epithelial state recovery dynamics. **E)** Time taken to regain 90% of epithelial phenotype share in the population on varying both the durations of EMT induction and S_02_. The results are presented for both with and without epigenetic regulation scenarios. Increasing period of EMT delayed epithelial state recovery for a given S_02_ values when epigenetic changes occur during EMT. The results presented were mean of three independent simulation runs of 10K cells each. Base parameters used for simulations were α = 0.15, S_01_ = 100K molecules, and S_02_ = 350K molecules, β_for_ = 240 hrs, and β_rev_ = 720 hrs, with specific ones being perturbed in their corresponding cases as is shown in the figure panel’s title or legend (A-E). The above parameters were assigned to every cell in the population. For cases without epigenetic regulation, α = 0.

**Fig S15.**
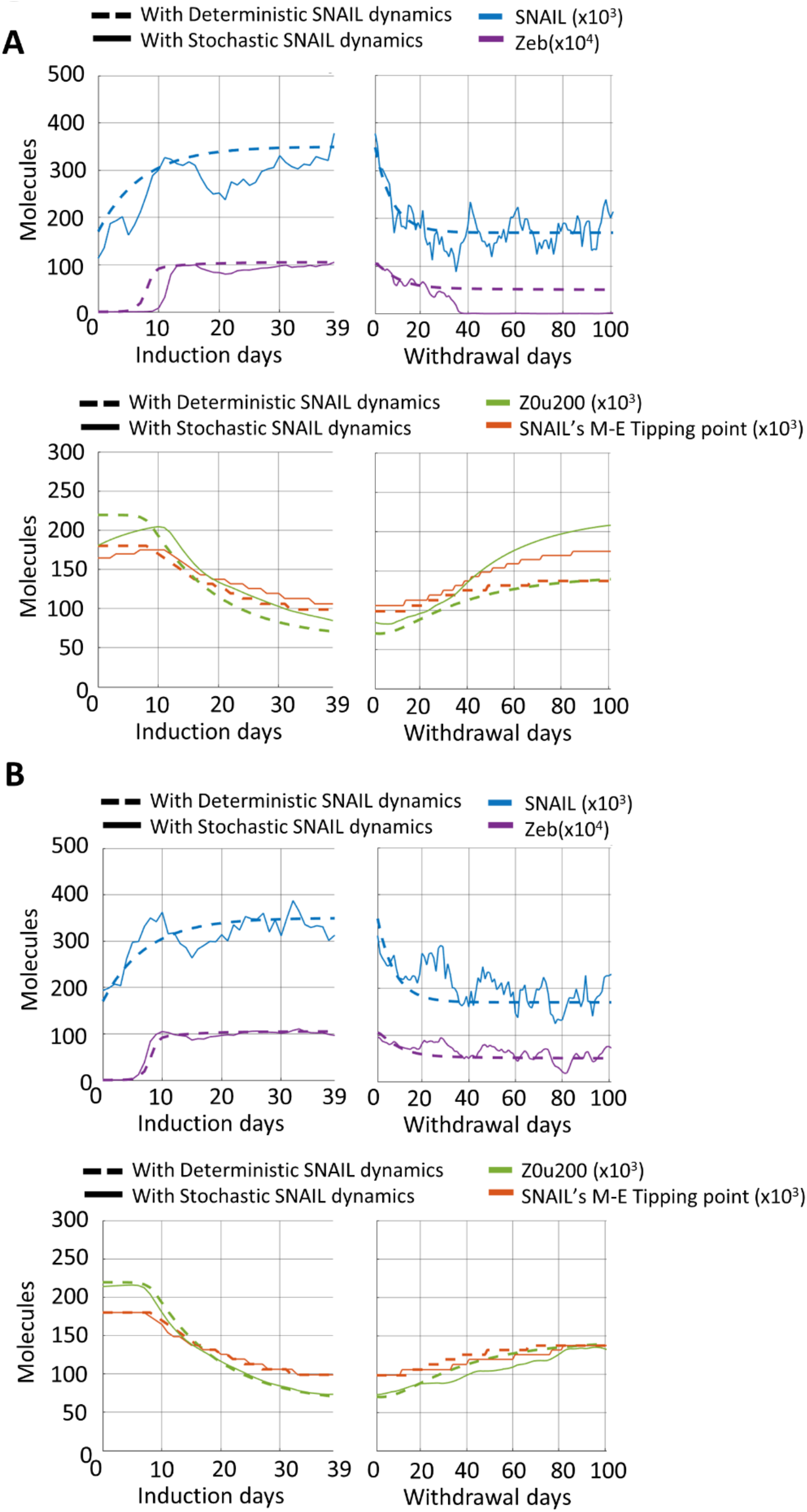
Stochastic SNAIL dynamics enables epithelial state recovery for higher S_01_ values. **A, B)** Stochasticity in SNAIL levels (solid curves) can cause MET of the cell during withdrawal period even for higher S_01_ where the cell with deterministic SNAIL dynamics (dashed curves) retained its M state. Plots generated using following parameters for simulations: α = 0.15, S_01_ = 170K molecules, S_02_ = 350K molecules, β_for_ = 240 hr, and β_rev_ = 720 hr.

**Fig S16.**
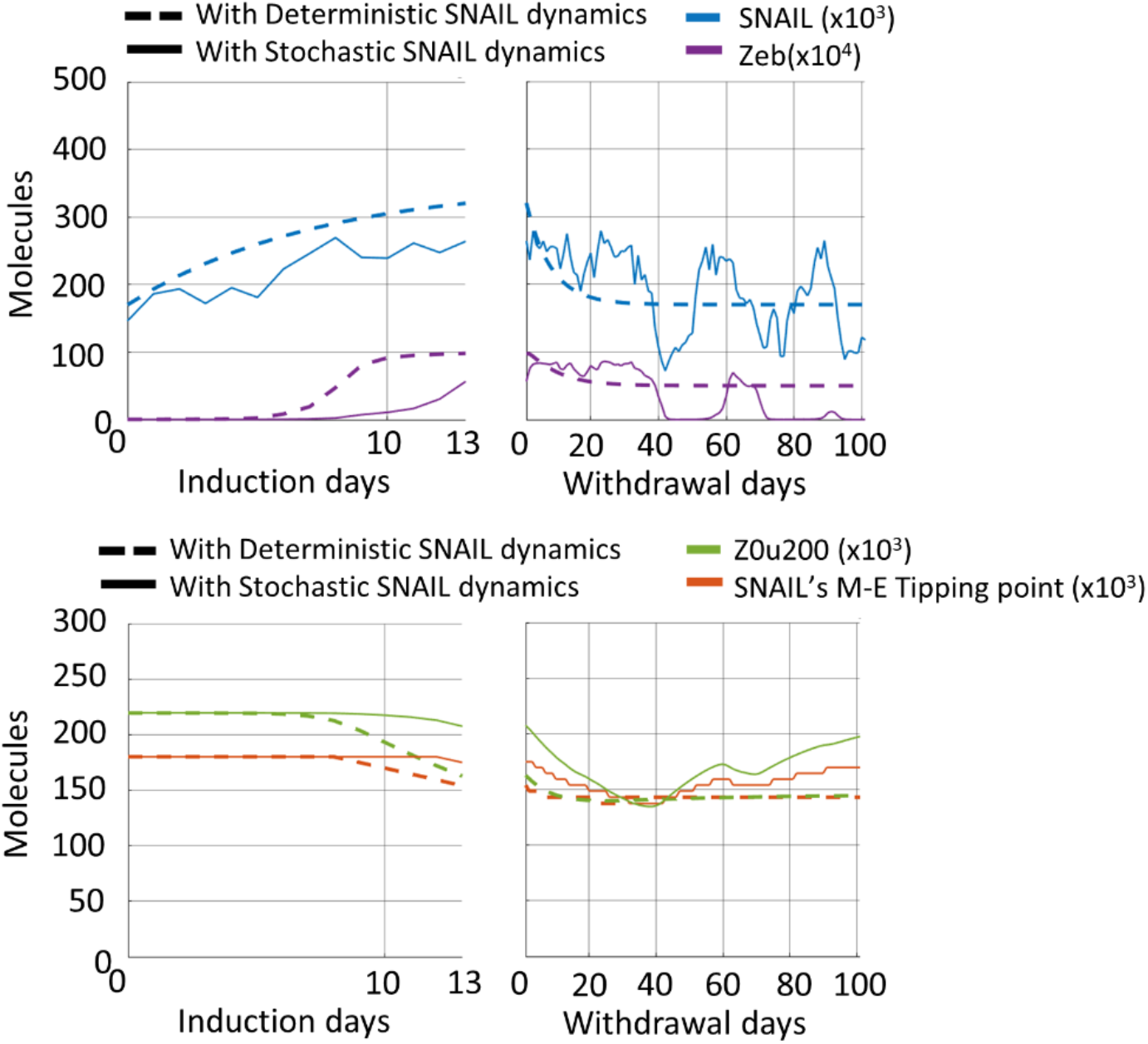
Stochastic SNAIL dynamics enabled plasticity in cell states for higher S_01_. Switching of mRNA ZEB levels between E and M phenotypic ranges were observed with stochastic SNAIL dynamics. Plots generated using following parameters for simulations: α = 0.15, S_01_ = 170K molecules, S_02_ = 350K molecules, β_for_ = 240 hr, and β_rev_ = 720 hr.

